# A Habitat Selection Multiverse Reveals Largely Consistent Results Despite a Multitude of Analysis Options

**DOI:** 10.1101/2024.06.19.599733

**Authors:** Benjamin Michael Marshall, Alexander Bradley Duthie

## Abstract

Researchers are intrinsically part of the research process. While we may strive for objectivity, there are always judgement calls required during research. When you ask ten researchers to answer the same question with the same dataset, you will likely receive ten different answers. This variation in answers has been linked to several disciplines’ replication crises. Here, we explore whether answers from movement ecology, specifically habitat selection, vary as a result of differing analytical choices. We conducted a multiverse analysis on around 400 synthetic animal movement datasets, exploring a multitude of analysis pathways to determine habitat selection, resulting in approximately half a million unique estimates of selection. By using simulated virtual animals with a known preference, we were able to show which decisions during analysis could lead to more variable estimates of habitat selection. The multiverse revealed that data quantity (i.e., tracking frequency and duration) was more important to obtaining consistent answers than any analysis choice. Overall, the pattern of estimates shows the majority of analysis pathways provide similar final results, particularly for modern analysis methods. The pattern reflects findings from other disciplines, indicating that while movement ecology is not immune to issues of non-replicability stemming from researcher choice, it is also not at any greater risk than other disciplines.

## 1 Introduction

Researchers are intrinsically part of the research process (Levins & Lewontin, 1985; Tang-Martínez, 2020), and our expectations can shape our conclusions (Holman et al., 2015). While we can strive to conduct research objectively, research commonly requires judgement calls (Steegen et al., 2016). Such choices occur throughout the research process, encompassing everything from study design (e.g., sample size, sampling intensity, sample stratification) to analysis (e.g., Bayesian or frequentist, handling of outliers). We draw on our own experience, and the input of peers, to try and ensure the best decisions are made to produce robust and reliable results, but we also contend with data, expertise, and interpretability constraints (Liu, Althoff & Heer, 2020).

Researchers are also influenced by the incentive system around them (Anderson et al., 2007; Receveur et al., 2024). We cannot undertake research in a vacuum; we often require institutions, funders, and scientific journals to produce and share research. These bodies can influence what research is conducted; they are in a position to incentivise or disincentivise research of certain topics or methodologies (Fanelli, 2010a; Ware & Munafò, 2015; Smaldino & McElreath, 2016). The use of impact factor (and similar metrics) is an example of citations being used as a measure of quality or research worth. But when examined closer, the impact factor appears detached from the robustness or reliability of the research (Brembs, 2018). A decrease in robustness can be seen in the increases in effect size inflation, p-value misreporting, and other measures of quality (Brembs, 2018). Similarly, novelty has been promoted by journals to the detriment of replication (Vinkers, Tijdink & Otte, 2015; Forstmeier, Wagenmakers & Parker, 2017; Brembs, 2019), despite widespread agreement on the importance of replications (Fraser et al., 2020). There is a bias towards positive, statistically significant results (Jennions & Møller, 2002; Cassey et al., 2004). Due to the nature of statistical significance, a prioritisation of significant results can elevate underpowered studies and boost false positive rates (Forstmeier & Schielzeth, 2011; Albers, 2019).

Unfortunately, there is evidence that the systems of incentives trickle down to impact the decisions of researchers while undertaking research and publishing results. The more detrimental of these decisions have been termed questionable research practices (Fraser et al., 2018; Bishop, 2019). Questionable research practices can be broadly viewed as methods to achieve a neat, statistically significant, and publishable narrative (O’Boyle, Banks & Gonzalez-Mulé, 2014). In the worst cases, narratives can be prioritised over transparently reporting results.

There is a fear that questionable research practices, and the broader incentives they are connected to, are responsible for the replicability crisis (also referred to as the reproducibility crisis). Across many disciplines, there are examples of replication studies being unable to replicate prior research (Freedman, Cockburn & Simcoe, 2015; Open Science Collaboration, 2015; Kelly, 2019). Often these replication efforts are conducted with larger sample sizes, or rely on the consolidation of many independent studies (often in the form of meta-analyses). The implication is not that the original studies were necessarily flawed; but, in the absence of questionable research practices, sufficient variation exists in the study subjects or methods to obscure a consistent effect (i.e., variation other than that stemming from sampling alone Simonsohn, 2015).

However, variation can also stem from analysis flexibility; i.e., the presence of many reasonable ways to analyse the same data to answer the same question. This flexibility helps enable questionable research practices by providing a suite of answers to choose between (Fraser et al., 2018), and is potentially steered by publication bias (Jennions & Møller, 2002; Cassey et al., 2004) if more publishable results are prioritised or rewarded over less exciting but robust results. Given the prominence of questionable research practices and publication bias, the inconsistencies between initial and replication studies warrant investigation (especially when analysis flexibility is also implicated in potentially flawed replications Bryan, Yeager & O’Brien, 2019). Even in the absence of any undesirable incentives, analysis flexibility can still lead to variable results simply as the result of researchers considering approaches of differing validities for a given dataset (Gelman & Loken, 2013; Gould et al., 2023).

Replications are a key tool for identifying potential sources of variation that lead to differing conclusions (e.g., subtle differences in methodologies or sampling). For some disciplines, such as ecology, replications can be difficult to justify due to the impacts on the study subjects and high monetary costs. Ecological systems are also complex and in constant flux, often frustrating perfect replications due to changes in space and time (Nakagawa & Parker, 2015; Schnitzer & Carson, 2016), leaving researchers with a distinct lack of direct replications in ecology (Kelly, 2019). The low prevalence of ecological replication studies makes it difficult to assess the overall rate of replication in ecology (Kelly, 2019), but there are several examples that suggest irreplicabilty is something ecologists should be wary of (Wang et al., 2018; Sánchez-Tójar et al., 2018; Roche et al., 2020; Clark et al., 2020). The potential for irreplicabilty is further supported by evidence of positive publication bias (Fanelli, 2010b, 2012), and links between smaller sample sizes and inflated effect sizes (Lemoine et al., 2016).

As direct replications can be difficult to conduct, ecology is often left to assess replicability via conceptual replications (Fraser et al., 2020) or efforts broadly referred to as quasi-replications (Palmer, 2000). Replications range in intensity. Direct (or exact) replications are attempts to replicate a tightly defined concept or hypothesis while duplicating all characteristics of the original study. Partial replications are a step looser, where the concept/hypothesis tested is less clearly defined (e.g., applicable to a broader scale), but efforts are made to repeat the same methodology. The most general category is conceptual replications, where the subject and method of study varies from the original study, but the replication targets a the same concept or hypothesis (Nakagawa & Parker, 2015; Kelly, 2019). Both partial and conceptual can be classed as quasi-replications if the concept and scale are broadly defined (Nakagawa & Parker, 2015).

Conceptual replications are valuable, but rely on researchers’ ability to compare replication efforts to previous findings. An important aspect of those comparisons is accounting for factors differing between the studies that are not salient to the effect of interest (Forstmeier, Wagenmakers & Parker, 2017), e.g., those linked to sampling differences (Simonsohn, 2015). An example of sampling differences leading to differences in results can be seen in the case of reptile space use. Silva et al. (2020) showed how the frequency at which a reptile was located (via radio-telemetry) by a researcher interacted with the space-use estimation method, leading to differences in area estimates even when using the same estimation method. This reveals how the choices during analysis (e.g., choice of area estimation method) impact results, and how the uncertainty introduced by those choices change depend on the sampling. It presents a scenario where the *correct* choice is dependent on preceding decisions; thereby, highlighting the need to explore the impacts of multiple decisions simultaneously.

As seen in the reptile space use example, the choices available to the researcher (i.e., researcher degrees of freedom, Simmons, Nelson & Simonsohn, 2011) is a key source of variation among studies. Research degrees of freedom (or flexibility in analysis, Forstmeier, Wagenmakers & Parker, 2017) have been elegantly demonstrated by a number of “many analysts” studies (e.g., Silberzahn et al., 2018; Huntington-Klein et al., 2021). In these studies, a number of researchers, or research groups, are tasked with answering the same question. Naturally each participant takes a slightly different approach, both in how the question is interpreted (Auspurg & Brüderl, 2021), and the analysis approach chosen (Gelman & Loken, 2013; Bastiaansen et al., 2020; Gould et al., 2023), resulting is different final results. Such variation in results can be sufficient to change the final conclusions (Salis, Lena & Lengagne, 2021), or at least alter the strength of an estimated effect (Desbureaux, 2021).

### 1.1 Multiverse analysis

An approach addressing the unknown impacts of undisclosed researcher degrees of freedom is to fully explore all plausible or reasonable analysis choices open to researchers, i.e., to explore a multiverse of design choices (Steegen et al., 2016). This multiverse analysis –closely linked to vibration of effects (Patel, Burford & Ioannidis, 2015), multi-model analysis (Young & Holsteen, 2017), and specification curve analysis (Simonsohn, Simmons & Nelson, 2020)– has the potential to quantify the variation stemming from researcher’s analyses choices (Rijnhart et al., 2021). Choices can include everything from from sample sizes and splits (e.g., Webb & Demeyere, 2021) to measurement and summary statistics (e.g., Parsons, 2020), but crucially should only include options that are reasonable (Simonsohn, Simmons & Nelson, 2020; Del Giudice & Gangestad, 2021). What counts as reasonable is not necessarily obvious, and inclusion of irrelevant choices can easily mask important choices because of the multiplicative nature of a branching path network (Del Giudice & Gangestad, 2021) (Fig. 1). Construction of a multiverse requires justification for which decisions are treated as variable, and why there is not an *a priori* and defensible single solution (Del Giudice & Gangestad, 2021). A multiverse populated with well-justified decisions allows the exploration of which choices inflate variation among analysis universes, while also offering insights into how to deflate variation (e.g., refinement of initial study design, the removal of ambiguities like tightening categories definitions, Steegen et al., 2016).

**Figure 1.**
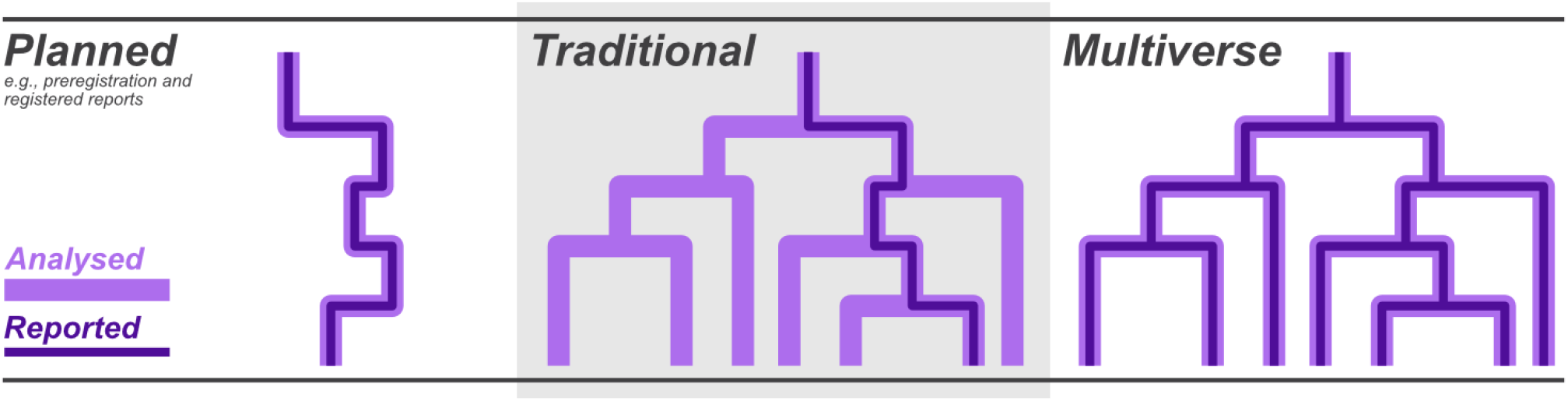
Diagram showing how multiverse analysis differs from other approaches. Each branch node represents a choice made during analysis. Based on Figure 1 from Dragicevic et al. (2019).

However, even if all choices are equally valid, multiverse analysis cannot simply provide a correct answer (Steegen et al., 2016). The “average” result is not necessarily the closest to the truth. If we were to undertake a multiverse analysis in a scenario with a “known truth”, i.e., using a simulated dataset (Bastiaansen et al., 2020), we may be able to identify the amount of variation attributable to different sources (e.g., biological variation vs study design variation, Breznau et al., 2021), and potentially the systematic biases stemming from specific choices. The use of simulated data is an established way to explore the robustness of methodologies (Minchin, 1987; Silva et al., 2020), and here we harness the benefits of simulated data to assess the impacts of researcher degrees of freedom on the results garnered from animal movement datasets.

As mentioned, ecological systems are complex to study and frustrate replication efforts (Nakagawa & Parker, 2015; Schnitzer & Carson, 2016); this can be especially true in the case of movement ecology and animal movement. Studying animal movement is often conducted using bio-telemetry/bio-logging devices that when attached to an animal record their location (Joo et al., 2022). From the resulting dataset of times and locations, researchers can derive a multitude of insights about the animal’s life, ranging from more surface, such as movement characteristics (e.g., speed, turn angle), to more in-depth such as home range or habitat selection. For the deeper insights a greater number of decisions are required during the analysis process, opening up the possibility that different researchers will take different pathways to answer the same question. With many forking pathways, analysis of movement data to determine an animal’s habitat selection presents a perfect setting for a multiverse exploration of how the researcher decisions impact the final results.

Here we construct a multiverse of analyses to explore how decisions concerning the design and analysis of an animal tracking data can impact the findings on habitat selection. Using a branching set of sampling and analysis options, we examine which decisions lead to the most variation in estimated habitat preference for three distinct simulated animal species.

## 2 Methods

### 2.1 Simulating the Scenarios

We simulated three scenarios each represented by a different species, on different landscapes. We simulated the landscapes using the NLMR v.1.1.1 package (Sciaini et al., 2018), and the animal movement using abmAnimalMovement v.0.1.3.0 package (Marshall & Duthie, 2022). The abmAnimalMovement simulations are a discrete time, agent-based modelling approach for simulating animal movement. Further details of how the scenarios were parametrised can be found in Marshall & Duthie (2022), and also in the abmAnimalMovement github repository along with associated documentation: https://github.com/BenMMarshall/abmAnimalMovement, also archived: https://doi.org/10.5281/zenodo.6951937.

The simulated species are as follows:

- Species 1, Badger: site fidelity to two shelter sites, a low movement speed constrained by terrestrial environment and territoriality, an 8-12 hour activity cycle with seasonal shifts.
- Species 2, Vulture: medium site fidelity via the use of multiple roosting/resting sites, a high and variable movement speed with minimal landscape resistance, an 8-12 hour activity cycle with seasonal shifts.
- Species 3, King Cobra: lower site fidelity making use of many shelter sites, a medium movement speed through a landscape with high resistance barriers, an 8-12 hour activity cycle combined with a approximately weekly forage-digest cycle and a weak seasonal cycle.

Each simulation generated a year’s worth of movement data with a data point recorded every minute, and we simulated 4 individuals per species. The full parametrisation of the simulated species can be found in the code associated with this paper, specifically the simulate_individual.R function (available at: https://github.com/BenMMarshall/ multiverseHabitat, and archived at: https://doi.org/10.5281/zenodo.12169335).

The agent-based animal movement simulations require an environment to guide movement. Our landscapes are defined as three stacked matrices, where the rows and columns of the matrix represent the X and Y locations in the landscape, and the value in each cell numerically describes the quality of a given resource the animal can consider when deciding where to move. Each of the three matrices’ describe one resource: foraging quality, shelter site quality, or movement ease (Fig. 2), which are evaluated by the animal differently depending on the behavioural state the animal is in (resting, exploring, foraging). All three are based on a single initial random generation using a Gaussian field.

**Figure 2.**
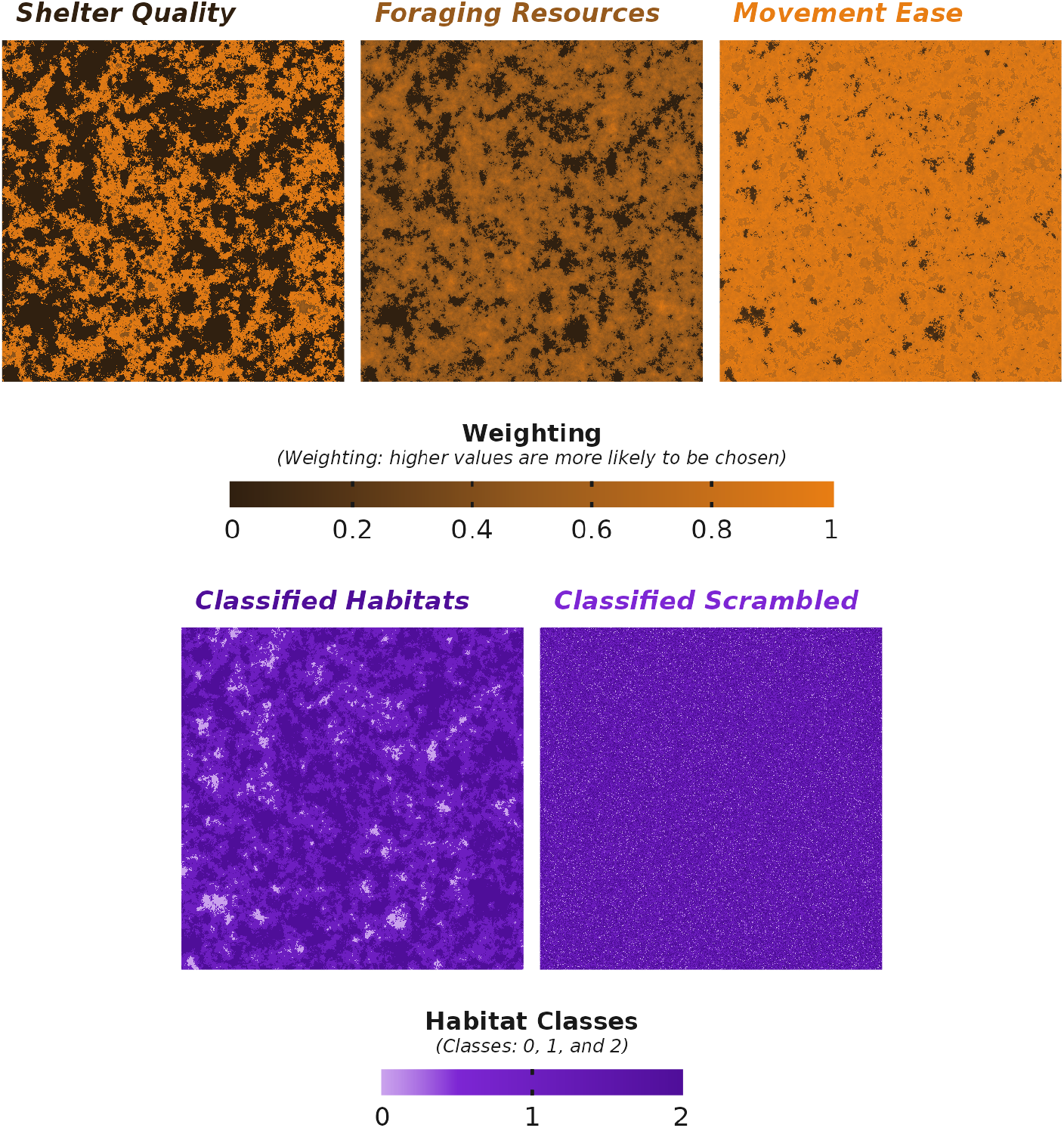
An example of the underlying environmental matricies used during the Badger species simulation, and the subsequent habitat classes to be used in the habitat selection analysis.

From the initial independent Gaussian fields, we altered the values to exaggerate the difference between high resource areas and low value areas; there-by generating non-independence of resources. Broadly, we created: core resource areas with higher foraging quality but lower movement ease, edge areas that overlap with the forest with better movement ease, and more barren areas with high movement ease but with minimal foraging value. All shelter sites occurred within the higher quality core resource areas.

While our simulation landscapes used to generate the simulated animal movement comprise of continuous values that affect the probability of a simulated animal using a cell or not, many habitat selection studies use categorical habitat metrics or land use types. Therefore, to simplify the interpretation of the multiverse, we simplified our landscape into three distinct categories for analysis (Fig. 2): a low quality area where resources are low but movement is easy (class 0 in the figure), an edge habitat with middling resources and high movement ease (class 1), and a high resource habitat with low movement ease (class 2). This results in a classified matrix/raster of habitat types usable for downstream analyses (Fig. 2). Class/habitat 2 was our target for all downstream analysis as it represents a preferred foraging area for the simulated species.

While the categorisation of landscapes is a major decision during the analysis of habitat selection, we elected to keep the landscape classification constant to maintain a feasible multiverse analysis. However, to provide a point of comparison and to begin to explore the impact of selection strength on the analysis choices and estimates, we re-ran the entire multiverse with a scrambled habitat raster. The scrambled habitat raster contained the same three classifications, but their positions were randomised, resulting in a habitat raster detached from the underlying simulated selection thereby representing low or no selection (i.e., random independent cell assignment, Fig. 2).

### 2.2 Sampling and Analysis Options

The number of variations and choices we were able to explore was ultimately dictated by computational costs and time. No multiverse can ever be entirely comprehensive, but we endeavoured to include the major decisions that are required during analysis and may not have immediately apparent single answers.

#### 2.2.1 Sampling

The first decisions concern data quantity. We varied tracking duration and tracking frequency, while keeping consistency fixed. While tracking consistency, or data loss during tracking, is an element affecting data quantity, the numerous ways of defining consistency led us to avoid exploring tracking consistency. Tracking timing is also an important consideration; for example, recording the locations of a diurnal animal only at night is highly unlikely to be a viable way of determining foraging choices. For this exploration, we assumed that the researcher has sufficient knowledge about the animal’s ecology to prioritise tracking during active hours. Timing only becomes a consideration when tracking frequency lowers to the point where the tracks performed during a day could occur entirely outside the animal’s active period. Therefore, for the lower tracking frequencies we ensured that daily tracking was centred around midday (as all our simulated species had diurnal activity patterns). We varied tracking duration from 7 to 240 days, and tracking frequency from 0.5 to 168 hours between subsequent data points (i.e., 2 to 0.01 points per hour).

Data or location accuracy is another aspect of sampling variation. Different bio-logging equipment, terrain, animal behaviour, and weather can all impact the location error when tracking an animal. The causes of and solutions to location error are numerous; therefore, we did not explore the impact of error, instead prioritising more variation in other decision nodes.

In summary, the sampling decisions covered: variation in three species represented by 4 independent individuals, tracking frequency, and tracking duration (Fig. 3).

**Figure 3.**
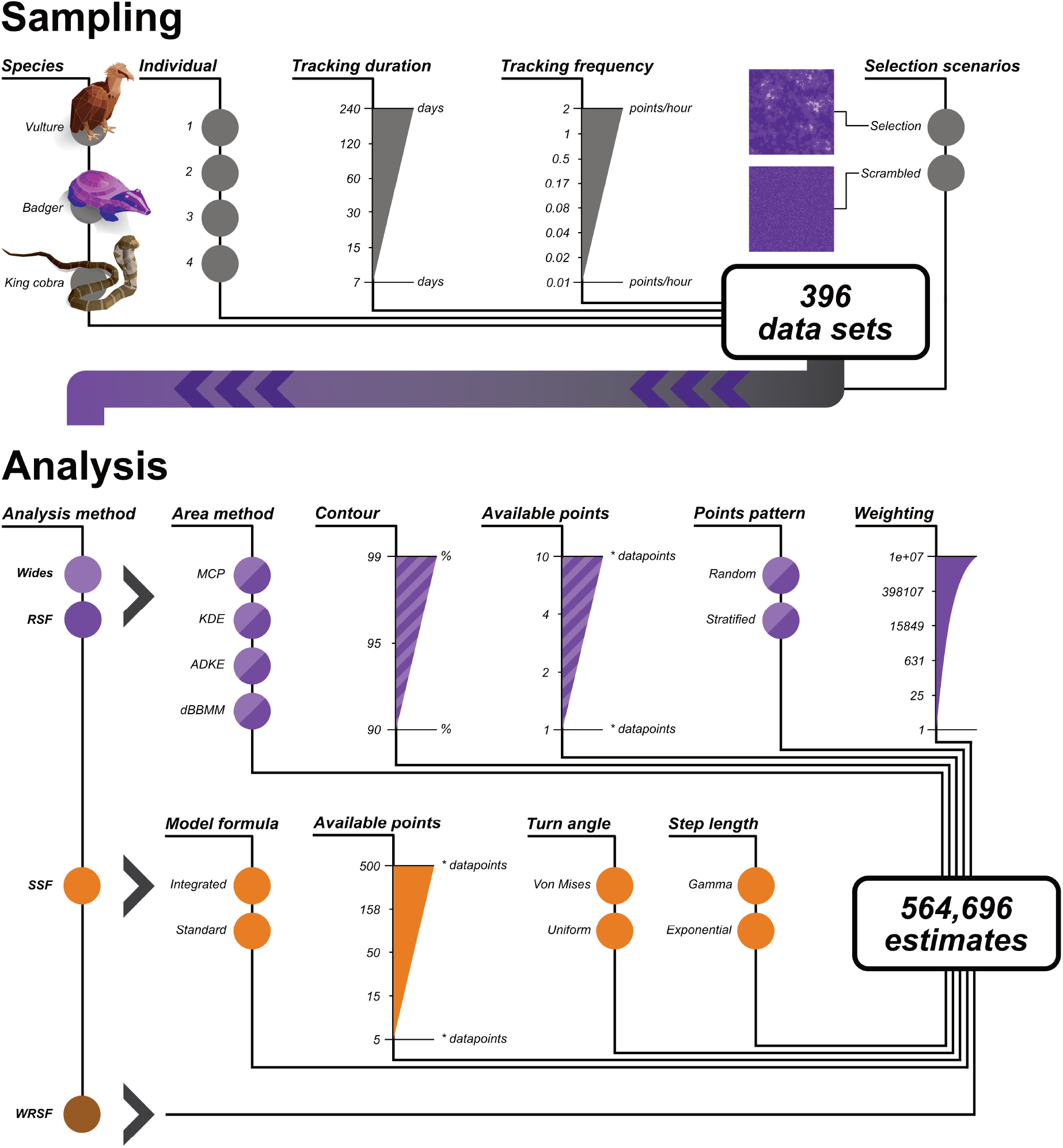
An illustration of the different decisions creating a multiverse examing animal movement. Grey choices pertain to data simulation and sampling. Species: the three different movement archetypes the simulations were loosely based upon. Individual: the number of individuals simulated per each species. Tracking duration: the length in days of the tracking data sampled from the overall simulated year. Tracking frequency: the frequency of the sampling of data points from the 1-per-minute simulated data. Selection scenarioes: the two classified habitat landscapes used during analysis. Purple choices concerned with analysis pathways using area-based estimations of availability (Wides: Wides selection ratios, RSF: Resource Selection Function). Area method: the method used to generate the available area polygon for the individual (MCP: Minimum Convex Polygon, KDE: Kernel Density Estimator, AKDE: Autocorrelated Kernel Density Estimator, dBBMM: dynamic Brownian Bridge Movement Model). Contour: the threshold used to convert the utilisation/occurrence distribution into a polygon. Available points: the number of generated points to extract available habitat within the polygon. Points pattern: whether the random points were generated in a stratified or random pattern. Weighting: the weighting of the available points in relation to the used points in the RSF formula. Light orange choices pertain to the SSF (Step Selection Function) analysis pathways. Model formula: whether the step and turn angles were added as interactions to the habitat variable (integrated if added). Available points: the number of randomly generated available points per step/used point. Turn angle: the distribution used to generate the turn angles of the available points. Step distribution: the distribution used to generate the step distances of the available points. Dark orange is the WRSF (Weighted Resource Selection Function) analysis. Decisions that required discrete choices are illustrated with circles, whereas those that continuous are depicted as triangles.

#### 2.2.2 Analysis

For the analysis, we focused entirely on decisions that result from an R analysis workflow (Fig. 3). R is the most common tool for analysing animal movement data (Joo et al., 2022). Joo et al. (2020) provide a review of the packages available to analysis movement data in R, and we used this review as a resource for determining the options for habitat analysis. Specifically, we explored a subset of decisions made during the workflows using adehabitatHS v.0.3.17 (Calenge & Mathieu Basille, 2023), and amt v.0.2.2.0 (Signer, Fieberg & Avgar, 2019) R packages. During the writing of this paper, a new method was described as part of the ctmm package v.1.2.0 (Fleming & Calabrese, 2023), which we also preliminarily investigated.

Combined, the packages offer many options for habitat analysis. We focus on four that tackle individual habitat selection:

- From the adehabitatHS package: resource selection ratios (Wides)
- From the amt package: Resource Selection Function (RSF)
- From the amt package: Step Selection Function (SSF)
- From the ctmm package: the newly described Weighted Resource Selection Function (wRSF Alston et al., 2023).

Each of the methods require downstream decisions resulting in a “garden of forking paths” to a final estimate of habitat selection (Fig. 3).

##### Area based approaches

The Wides and RSF methods share many analysis decisions. The Wides approach aims to calculate selection ratios of used habitat compared to those available (Manly et al., 2007); whereas the RSF approach uses a logistic regression to compare the used and available habitats. Therefore, a principal concern in both approaches is how available habitat is defined.

The first decisions is whether to approach the habitat selection analysis as a type I (population habitat use compared to landscape-level habitat availability), II (individual habitat use compared to landscape-level habitat availability) or III design (individual habitat use compared to individual-level habitat availability). We are concerned with an individuals’ habitat selection, so ignored type I. While type II habitat selection can identify individual selection, it requires habitat availability to be defined on a population or landscape scale (i.e., a universal availability). As movement data presents an opportunity to estimate individual selection, and we are not simulating a true interacting population, we focus on type III. Type III and the need to individually define available habitat opens up many different analytically decisions concerning how that availability is defined.

We used a range of methods to build areas that could be considered available to the animal from the recorded locations of the animal: Minimum Convex Polygons (MCPs), Kernel Density Estimations (KDEs), Autocorrelated Kernel Density Estimations (AKDEs), and dynamic Brownian Bridge Movement Models (dBBMMs).

Minimum convex polygons (MCPs) simply creates a polygon based on the outermost locations, where the interior angles sum to less than 180 degrees.

Kernel Density Estimations (KDEs) use kernel smoothing to build a heat map of use or utilisation. Critical to KDEs, operation is a bandwidth or smoothing factor (h) that alters the resulting area treated as used by the animal (Silva et al., 2020). For this analysis, we use the reference bandwidth (href) as it is commonly used (Crane et al., 2020), while also quickly and consistently calculable on a individual-by-individual basis. An alternative common method to determine smoothing factor is Least Squares Cross Validation (LSCV). However, we avoided the LSCV method for two reasons: 1) the LSCV method can fail to converge on a smoothing factor, thereby introducing an unpredictable fail state in any potential multiverse, and 2) the more limited area estimate it produces makes it less suitable for defining habitat availability.

The limitations of MCPs and KDEs have prompted the development of newer methods of area use estimation (Silva et al., 2020) that better account for the non-independence and autocorrelative structures within animal movement data (Noonan et al., 2019). We used two of these methods to define availability.

First is the Autocorrelated Kernel Density Estimations (AKDEs) (Fleming et al., 2015; Fleming & Calabrese, 2017) from the ctmm package (Calabrese, Fleming & Gurarie, 2016; Fleming & Calabrese, 2023). The ctmm package provides a workflow for creating an area estimate based on a number of movement models. The movement models include different levels of autocorrelative structure: Ornstein-Uhlenbeck (OU) models account for central tendency; the Ornstein–Uhlenbeck Foraging (OUF) models build on this by also accounting for autocorrelation in movement speed; while the Independent Identically Distributed (IID) models assume location independence, meaning they are similar to traditional kernel density approaches. Each model type is also fitted in isotropic (circular) or anisotropic (elliptical) forms. The movement model used would constitute a decision during analysis; however, the ctmm package allows for model comparisons (using AICc) to chose a single best model. Therefore, we use the guidance from (Silva et al., 2022) to generate weighted AKDEs using the perturbative hybrid residual maximum likelihood method (pHREML), and select the best performing by AICc for inclusion in further analysis. Using weighted AKDEs means the area estimates better address gaps between locations (Silva et al., 2022), with no apparent drawbacks nor prohibitively large computational cost.

The second movement-specific method we used was dynamic Brownian Bridge Movement Models (dBBMMs) (Kranstauber et al., 2012) from the move package (Kranstauber, Smolla & Scharf, 2023), which estimates movement capacity of the animal to calibrate repeated random walks between known locations. DBBMMs require a window and margin size that defines the number of data points over which movement capacity (motion variance) is calculated (Kranstauber et al., 2012). Window and margin are defined by a number of data points; therefore, to keep the time they represent the same between different tracking frequencies, we changed their values for each tracking frequency. As our most infrequent tracking is 168 hours (7 weeks), we set the window to the number of data points collected over 168 hours, and a margin of 48 hours. The broader window and margin sizes helped reduce computational costs, a consideration when many dBBMMs require calculation.

Unlike the AKDEs, dBBMMs do not produce utilisation distributions. As dBBMMs are estimating the uncertainty surrounding potential movement pathways, the resulting distribution is better described as an occurrence distribution. This occurrence distribution describes the within-sample uncertainty rather than the potential areas available to the animal beyond the sampling period (i.e., it possess little to no predictive capabilities). Therefore, using dBBMMs to define habitat availability means rather than comparing the habitat use (derived from the movement data) to availability (broader area predicted by the movement data), we are more comparing habitat use (derived from the movement data) to an alternative measure of habitat use (an area estimating uncertainty surrounding the movement pathways). We included dBBMMs to examine whether this arguably easy-to-make conceptual mistake impacts the final results (Alston et al., 2022).

What is considered available to an animal is not cleanly translated from a biological concept to a statistical one. While the area estimation methods (MCPs, KDEs, AKDEs, dBBMMs) described above make –to varying degrees– use of the movement information contained within the dataset, they remain abstractions of the underlying drivers of animal movement (i.e., selection and perception). Therefore, the researcher must make a somewhat arbitrary choice regarding what is included/excluded and counted as available; each of these area estimation methods requires a choice to be made regarding the outermost boundary. We explored the impact of using a 90, 95, and 99% contour for all the methods. MCPs areas are generated based a percentage of the location points, whereas the KDE, AKDE, and dBBMM methods require a contour to be extracted from the utilisation distribution (occurrence distribution for dBBMMs). AKDEs also provide a 95% confidence interval surrounding any chosen contour; for the purposes of simplicity, we only use the point estimate for each % contour.

Once we had created a defined availability area, we generated points at which the habitat type is recorded to estimate the relative availability. How many, and in what alignment, these availability points are generated poses two additional points of analytical choice. For this study, we varied the number of availability points as a multiple of the number of animal locations in the dataset (from 1 to 10, Fig. 3). For the alignment of the points, points were either random or stratified within the available area previously defined.

As mentioned, resource selection functions (RSF) share the above decisions on availability with the Wides methods. In addition, for the RSF we also explored the impact of varying the weighting of the available points when the RSF (i.e., generalized linear model / logistic regression) model is run, which impacts the fitting process. There are suggestions that altering the weighing can improve model convergence and decrease the uncertainty surrounding habitat selection estimates (Fieberg et al., 2021); we included this decision to examine whether the pursuit of a more confident answer is biasing the point estimate (from 1 to 1e+07).

##### Step selection functions

In addition to the area based methods of Wides and RSF, we explored of Step Selection Functions (SSF) and Weighted Resource Selection Functions (wRSF).

Instead of estimating habitat availability using area, SSFs use observed step lengths and turn angles to generate available locations at each time step (in our case for each data point in the dataset; strata). Consequently, SSFs have a number of decisions not shared with the area methods.

We explored three decisions associated with the generation of random (available but unused) locations for each step. The first is similar to the area methods; we varied the number of random locations generated per step from 5 to 500. The second and third decisions concern the distributions used to generate the random step and turn angles. Step lengths were tested with a Gamma and Exponential distributions, while turn angle was tested with Von-Mises and Uniform distributions.

The other decisions we explored in SSFs were whether to run the model as a standard step selection or an integrated step selection function. The standard step selection model is a conditional logistic regression, where we aim to identify the association between whether a location is used/unused and the habitat values at that location (case_∼ values + strata(step_id_)). The integrated step selection model is very similar, but the predictors also included step length and turn angle as interactive terms with the habitat values. The addition of these two components is meant to reduce bias by better accounting for the selection that may simply be an artefact of the movements of the animal rather than active habitat selection (Avgar et al., 2016).

We have neglected to explore an expansive aspect of SSFs regarding high resolution data and the use of “bursts”. Bursts are used when the researcher wants to look at selection over a given timeframe; therefore, locations are grouped into bursts and the bursts become the strata in the model rather than timestep. Such grouping of data deserves investigation, but there are myriad of ways bursts can be defined, and their interaction with tracking duration, frequency, and consistency would warrant a separate multiverse analysis. As such, we have restricted this exploration to step selection functions where the datapoints are equal to the strata in the model.

##### Weighted Resource Selection Functions

Compared to the other methods, Weighted Resource Selection Functions do not have many associated choices. Part of the motivation behind their development was to integrate autocorrelation and other critical structures of movement data into the modelling of habitat selection (Alston et al., 2023). This contrasts to the other methods that either ignore the confounding effect of these structures, or rely on potentially arbitrary decisions for how to mitigate any biasing effects the structures produce. As a result, we have no exploration of analysis decisions for this method because aspects such an available area or points no longer apply, and availability is estimated as part of the model (Alston et al., 2023). Instead we explore only variations in data quantity (i.e., tracking duration and frequency).

### 2.3 Assessing the multiverse

To generate the multiverse, we used a meta-programming approach via the targets v.1.6.0 and tarchetypes v.0.9.0 R packages (Landau, 2021a,b). The targets package was created to facilitate reproducible workflows, but is also paired with functionality enabling multiple analysis routes and parallel processing. Targets’ ability to track analysis task progress and intermediate R objects reduces repeat computation in a complex workflow that may not be able to be run in a single sitting. We created a branching analysis workflow based on the decisions we wished to explore, then used targets to iterate over all combinations of all decisions and compile all analysis end points ready for further examination. We set a seed during the running of the multiverse to further aid reproduction of these results. The single seed means that all simulated individuals, and subsequent analysis using randomness are unique within the multiverse.

Further examination was complicated by the four methods described above producing habitat selection values on different scales with different decisions associated with them. We elected to analyse the impact of the decisions separately. For each method, we ran two Bayesian regression models to explore: 1) which decisions best explain the absolute deviation from the median estimated selection, 2) which decisions best explain the raw deviation from the median estimated selection. The first model provides us with an idea of the decisions that can lead to random, but potentially unbiased variation surrounding the median estimate; whereas the second model highlights decisions that lead to systematic over or under estimation of a habitat selection. All models included individual and species as nested group effects, as well as a group effect accounting for whether the classified landscape was scrambled or not. For example, the formula for the SSF model looking at absolute deviation: “absolute delta from median estimate”∼ 1 + “tracking duration scaled” + “tracking frequency scaled” + “integrated formula” + “step distribution” + “turn distribution” + “available points per step scaled” + “(1|landscape scrambled)” + “(1|species/individual)”. We normalised all predictor variables because they all existed on vastly different scales, as we wanted to assess relative impact. We assessed Bayesian model convergence using R-hat values, acf, and trace plots. Based on these assessments we modified the running parameters (RSF 500 iterations, 150 warmup; Wides 1500 iterations, 500 warmup; SSF 1500 iterations, 250 warmup; wRSF 2000 iterations, 500 warmup;).

We elected to have the model try and predict the deviation from the median estimate of habitat selection because the simulation cannot provide a direct analogous value to the outputs of all four methods. The analysis closest to mirroring the agent-based model of the simulated data is the SSF. The simulation is a discrete time model, where the animal makes decisions based upon a number of options making it similar to a SSF step-wise model structure. Despite this similarity, the decision-making of the animal is made on two time frames; thereby, breaking our ability to directly compare simulation values with step selection outputs. Additionally, the simulated selection of the animal is balanced against other demands such as site fidelity, movement resistance, and avoidance of certain locations; therefore, any input habitat selection into the simulation becomes directly incomparable with a derived answer (i.e., analysed) from the simulated movement. The closest analogous values we could generate came from running RSF and iSSF models directly on the simulated decisions made by the animal. We ran these models for both time scales: movement decisions every time step, and destination decisions every behavioural state switch. These model results, while not strictly comparable to the other outputs, help confirm the animal was correctly preferring the chosen habitat. The estimates calculated this way likely underestimate selection further as the rest sites were only generated within the preferred habitat (a simulated strict selection). The lack of direct comparability between simulated parameters and analysed estimates of selection supports the use of the median estimate to explore the cause behind the most deviant estimates.

## 3 Results

The completion of the multiverse resulted in 396 datasets representing an individual animal’s movements sampled via a distinct tracking regime (i.e., a combination of tracking frequency and duration). The datasets covered 3 virtual species, 4 individuals per species, frequencies from 0.006 to 2 locations per hour, and durations from 7 to 240 days. Each of these datasets underwent various combinations of analysis choices resulting in 564,696 estimates of habitat selection (Wides: 76,032; RSF: 456,192; SSF: 31,680; wRSF: 792).

### 3.1 General patterns

We can get an initial impression of the pattern of the final selection estimates using a specification curve (Simonsohn, Simmons & Nelson, 2020; Hall et al., 2022). In a scenario where the decisions make little difference, all estimates should group together around a common answer, with the majority of deviation from that common answer being the result of sampling or individual variation.

Starting with the Wides results, we see a quite gentle *S* shape (Fig. 4), with most estimates falling in the range of 0.5 to 2. There is an agreement of positive selection for habitat 2 as expected, and a stronger selection identified in the selection scenario compared to the scrambled raster (no selection) scenario. While the positive selection appears in reasonable agreement that selection is occurring (more estimates toward the higher values than lower), the exact estimate appears relatively sensitive to the choices made. Looking at the specific choices, we can see several weak patterns. The first is in tracking duration where we see a slight reduction in the spread of estimates with longer tracking durations, when selection is present. The same is not clear in tracking frequency because the lower tracking frequency could not be combined with lower tracking durations resulting in fewer estimates for low frequency scenarios. Despite this, we do see a slight reduction in estimate variation from frequencies of 0.17 locations per hour and higher, as well as more consistent positive selection estimates. Similar to the tracking intensity choices, the contour used during the area generation appears to affect the spread of estimates, with a larger contour leading to a slightly greater agreement between estimates. So too does the multiplier of available points, with more points slightly reducing the spread of estimates, although we see no movement in the median estimate. Higher numbers of available points, and a higher tracking intensity, did appear to lead to a greater chance of a successful estimate being produced (see numbers to the right of the 0 line in Fig. 4). The choice of area method (MCP = Minimum Convex Polygon, AKDE = Autocorrelated Kernel Density Estimate based on the best performing movement model, KDEhref = Kernel Density Estimate using reference bandwidth selection, dBBMM = dynamic Brownian Bridge Movement Model) appears to suggest more consistently similar estimates with AKDE and KDEhref methods compared to MCPs and dBBMMs, with MCPs leading to slightly lower estimates in the selection scenario. This difference disappears under the scrambled habitat classification, and in that scenario all analytical choices appear to make little impact on the median estimate.

**Figure 4.**
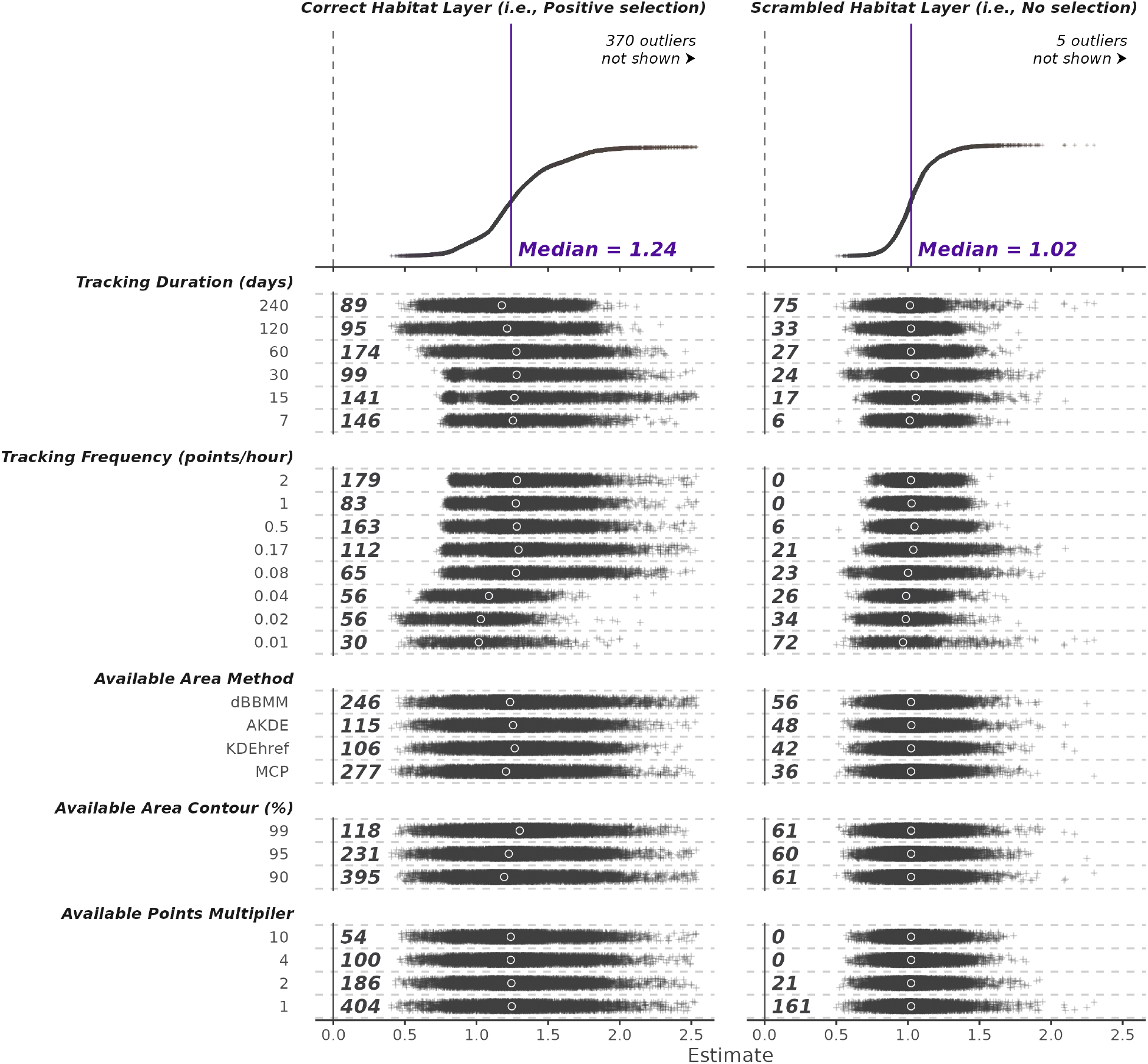
A specification curve showing all habitat selection estimates from the Wides analysis pathways applied to the simulated data. Each point is a estimate of habitat selection. Top plot shows all estimates according to estimate, with an arbitary sorted y-position. Lower plot show the same estimates in relation to the decisions that lead to them. Left-hand numbers describe the number of NA results connected to each decision, indicating a failure somewhere in the calculations. Solid circles are the medians for each choice.

The Resource Selection Function (RSF) specification curves shows a more ideal *S* shape, with the majority of estimates clearly converging on a common answer (Fig. 5). Similar to the Wides analysis, the majority of the answers and medians show selection for habitat 2 in the selection scenario. The no selection scenario still overall shows selection, but it is considerably lower. While there are answers incorrectly suggesting avoidance of habitat 2, all bar two medians connected to a specific choice (the lowest two tracking frequencies) suggest selection. To best see the changes in variation connected to a given choice, we can look at the rates and position of the outliers (highlighted in purple and orange). Although weak patterns, tracking frequency, and multiplier of available points see slight reductions in the number of estimates very high or very low. This is particularly apparent in the no selection scenario, where higher tracking frequencies led to very similar answers surrounding zero. Other patterns are difficult to discern, except for the weighting of used points that has a clear impact on the outlying low estimates. The distinct step-like pattern is a result of the logarithmic scale the weighting varies over (required to capture such a wide range of possible values), which can been seen reflected in other choices’ groupings of estimates. The weighting does not appear to impact the overall median, with weighting only appearing to dramatically impact the outlying results. Comparing the selection scenario to the no selection scenario, we can see that the selection scenario is more likely to produce larger outlying results, both positive and negative.

**Figure 5.**
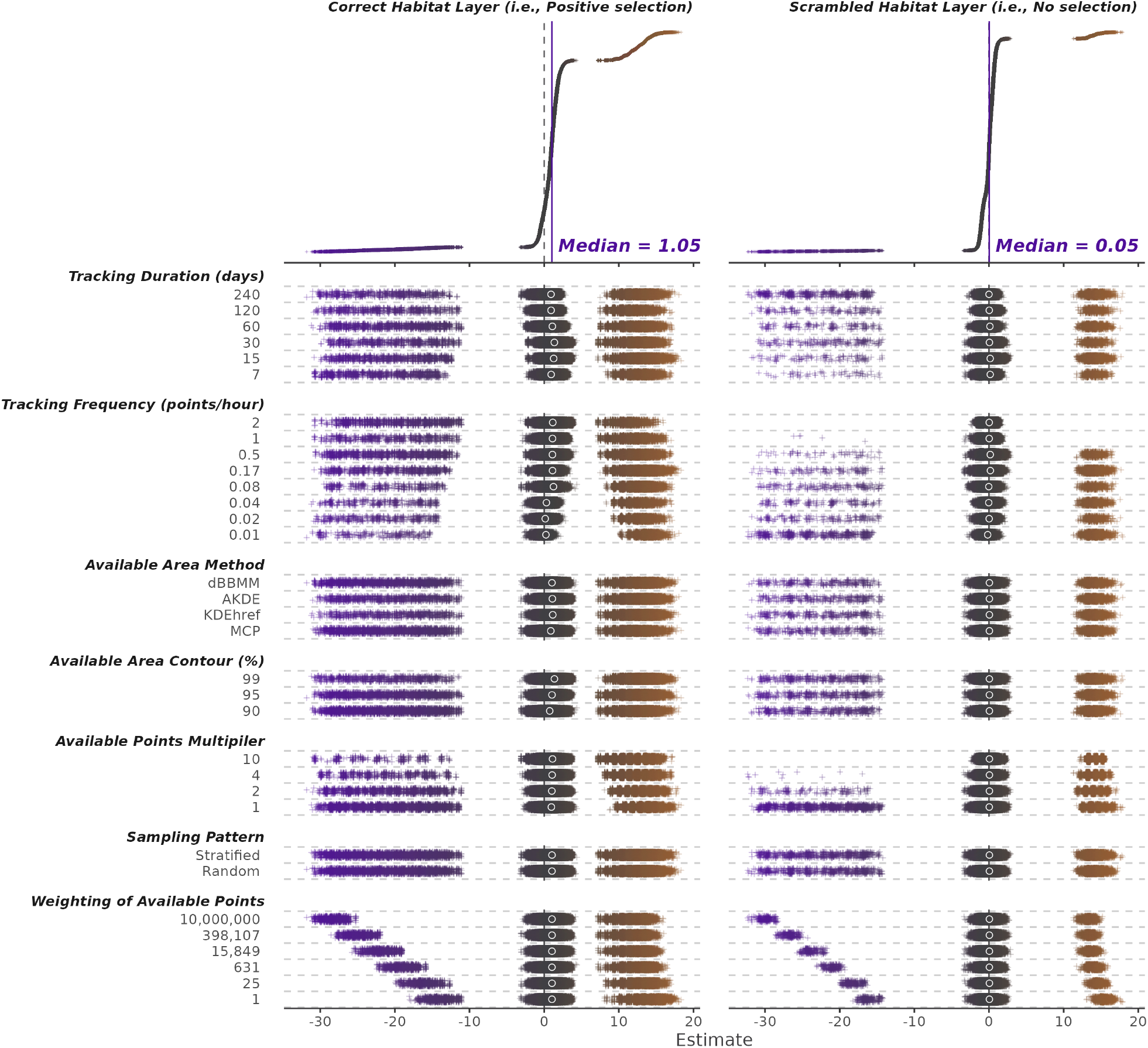
A specification curve showing all habitat selection estimates from the RSF analysis pathways applied to the simulated data. Each point is a estimate of habitat selection. Top plot shows all estimates according to estimate, with an arbitary sorted y-position. Lower plot show the same estimates in relation to the decisions that lead to them. Solid circles are the medians for each choice. Colours illustrate deviation from the median.

For the RSF analyses, we also recovered the range of the 95% confidence intervals surrounding each estimate. When plotted against the estimate, we see clear groupings where those most extreme estimates are also, by a significant margin, the most uncertain (Fig. 6).

**Figure 6.**
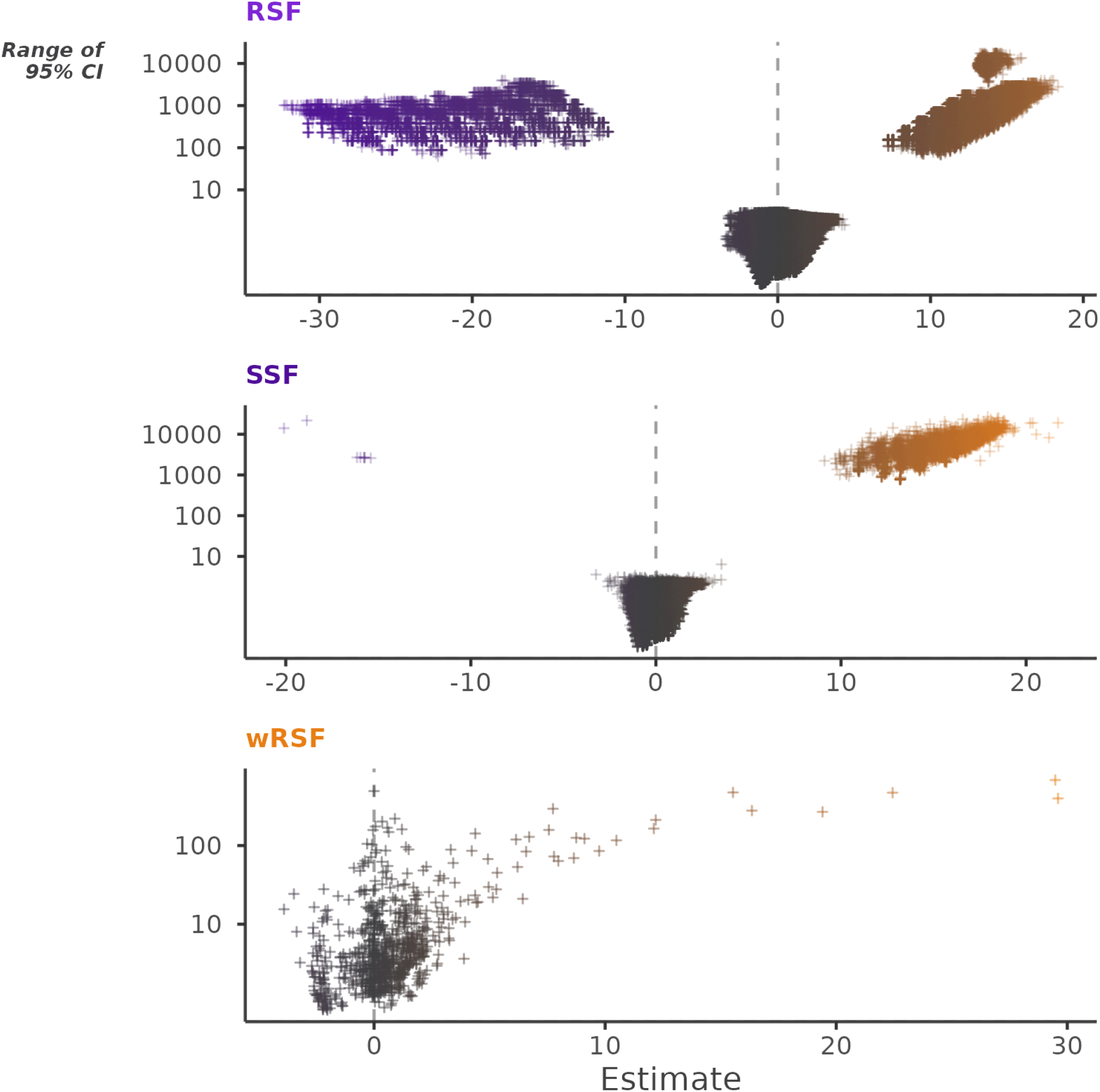
All standard errors associated with estimates of habitat selection from RSF, SSF, and wRSF analysis pathways. More extreme estimates of selection are associated with dramatically larger confidence intervals, N.b., y axis is log scaled.

The Step Selection Function (SSF) specification curves appears even more successful than the RSF. There is a reasonably clear *S* shape indicating many analysis end points resulting in similar estimates of selection (Fig. 7). There are also considerably fewer estimates that indicate avoidance of habitat 2. The stand out patterns are more easily seen in the outlying high estimates. Once again, there are indications of fewer outlying high estimates when data quantity increases, both with duration and frequency. The number of points per step (or strata) appears to intuitively follow that more available steps leads to more consistent estimations, and also the outliers that exist appear to become less extreme. The decisions concerning the distributions used to model step lengths and turn angles, as well as whether the model is integrated or not, appear to make little difference to the spread of final estimates. The impact of analysis choice on the SSF method appears to be more consistent between the two selection scenarios.

**Figure 7.**
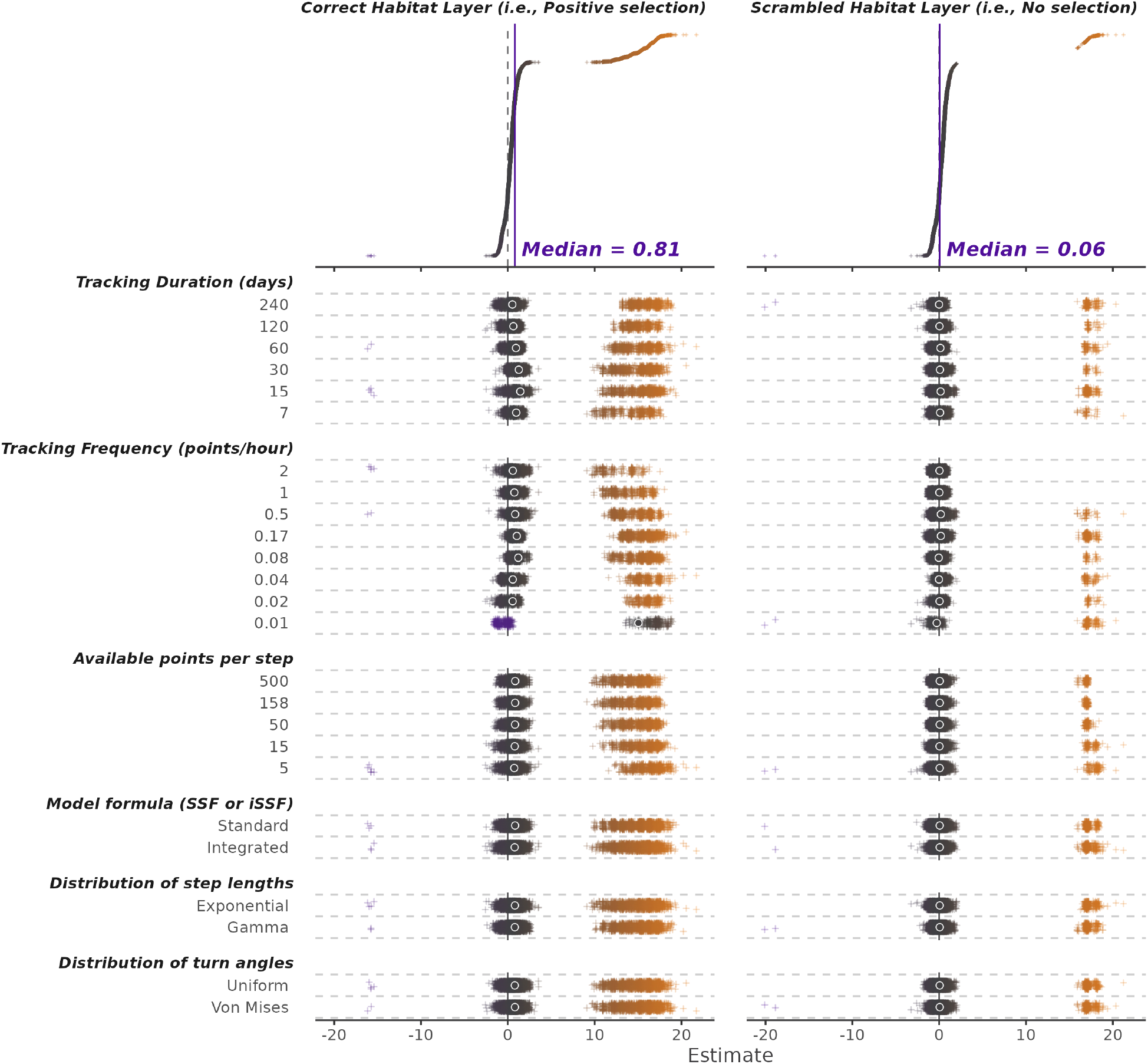
A specification curve showing all habitat selection estimates from the SSF analysis pathways applied to the simulated data. Each point is a estimate of habitat selection. Top plot shows all estimates according to estimate, with an arbitary sorted y-position. Lower plot shows the same estimates in relation to the decisions that lead to them. Solid circles are the medians for each choice. Colours illustrate deviation from the median.

Very much like the RSF explorations of the raw data, the 95% confidence intervals are dramatically larger (between 100 and 1000 fold) for the outlying estimates than those nearer the median estimate (Fig. 6).

The Weighted Resource Selection Function approach has very few decisions associated with it currently; however, we still explored the impact of tracking duration and frequency on the estimates (Fig. 8). The highest estimates appear connected to the lower tracking frequencies, but like the SSF results, those high estimates also tend to have large confidence intervals (Fig. 6). The bulk of the estimates agree and are tightly concentrated around the median with very few underestimating. As with RSF and SSF methods, the wRSF is very likely to correctly identify the positive selection and the median selection under the no selection scenario is lower, particularly when more data are available.

**Figure 8.**
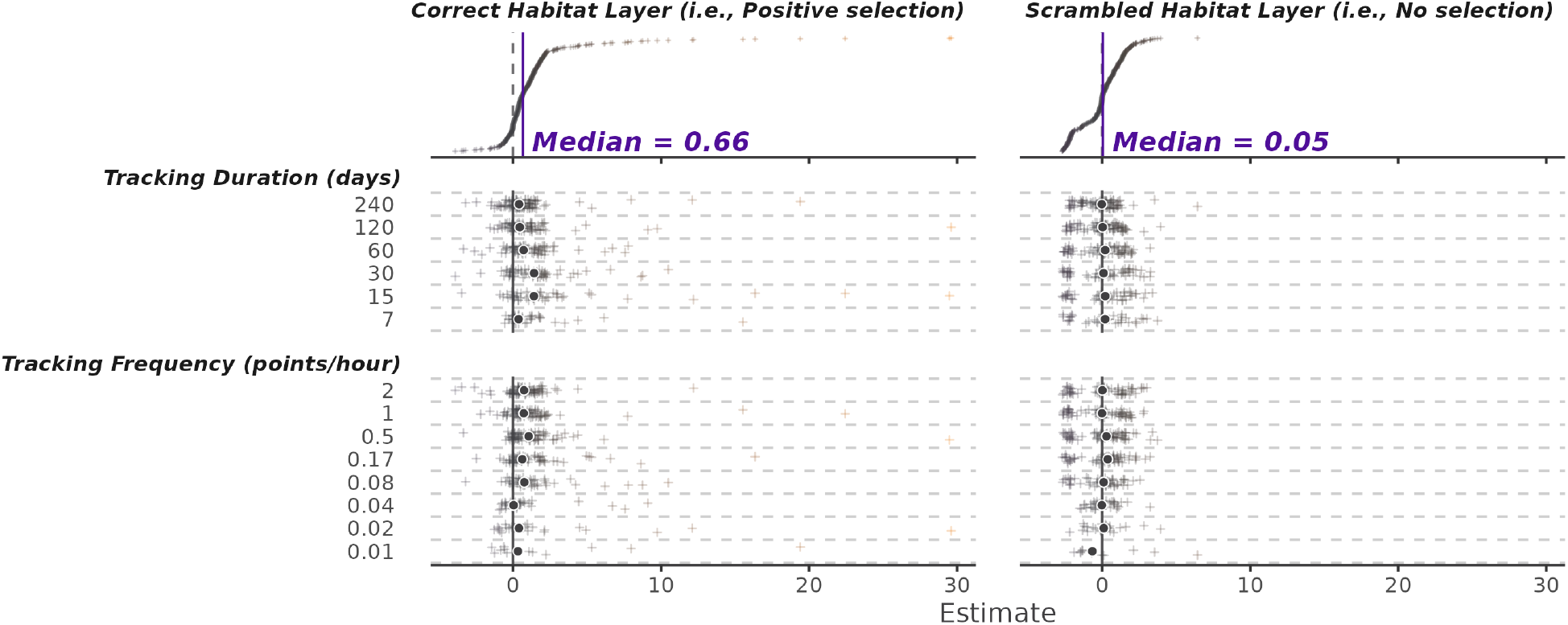
A specification curve showing all habitat selection estimates from the wRSF analysis pathways applied to the simulated data. Each point is a estimate of habitat selection. Top plot shows all estimates according to estimate, with an arbitary sorted y-position. Lower plot show the same estimates in relation to the decisions that lead to them. Solid circles are the medians for each choice. Colours illustrate deviation from the median.

### 3.2 Model exploration

While the specification curves are a good way of getting an intuitive sense of variation, the scale and range of estimates can skew the interpretation of which choices have the largest impact. Our Bayesian models cut through the noise displayed in specification curves to reveal more generalised patterns and absorb variation stemming from the stochastic nature of the animal simulations and the variation between species.

All models showed that the variation stemming from the analysis choices was less than that explained by the group effects of individual, species, and whether the landscape classification was scrambled (Table. S1). The conditional *R^2^* values ranged from 0.08 to 0.3, with the RSF models performing the worst. Largely the marginal *R^2^* values were very low (often below 0.1), meaning that while the models reveal clear effects, the overall variation they ascribe to given analysis choices is reasonably limited compared to the overall variation stemming from the different species, individuals, landscape classification, and stochastic nature of the movement data simulation.

For the decisions connected to Wides results, the absolute deviation was curtailed most effectively by increases in data quantity (Fig. 9; Fig. S1), with increases in tracking duration (*β*: −0.024, 95% CrI −0.026 – −0.021) being marginally more effective than tracking frequency (*β*: −0.027, 95% CrI −0.029 – −0.024). Of the other decisions, only the multiplier of available points appeared to decrease absolute deviation unambiguously (*β*: −0.004, 95% CrI −0.005 – −0.003), and its effect was much smaller than those concerning data quantity. Choice of area method as well as the pattern used to generate random points seem to make little clear difference, with all credible intervals connected to those choices overlapping zero. Contour choice increases the absolute deviation from the median estimate *β*: −0.004, 95% CrI −0.006 – −0.003. That appears counter to the initial specification curve plot and highlights the difficulty of identifying patterns from the raw data alone.

**Figure 9.**
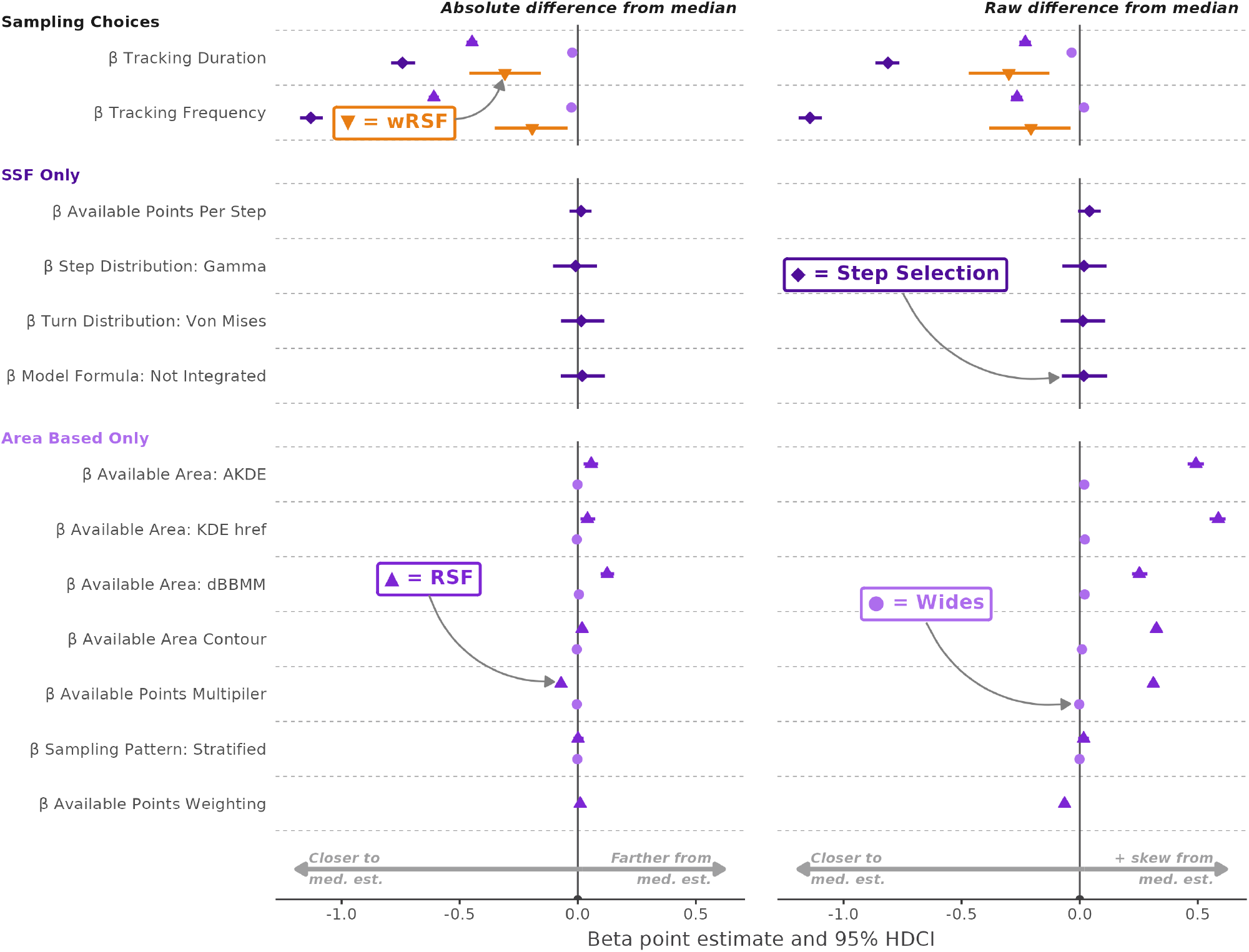
Estimated betas for each choice’s impact on the deviation from the median habitat selection estimate. Deviation from the median estimate was examined as the absolute deviation (left) and the raw deviation (right). All eight independent models are displayed together, split between analysis method and deviation type. Each point represents the point estimate derived from the 95% median highest density continuous interval, and is plotted alongside a line showing the extent of the 95% credible interval. The y-axis shows all the choices explored in the different analysis methods, grouped by the methods they apply to.

The model examining raw deviation from the median estimate reveals that the Wides approach is contrastingly impacted by tracking frequency *β*: 0.018, 95% CrI 0.015 – 0.021 and tracking duration *β*: −0.034, 95% CrI −0.037 – −0.031, with the former leading to greater estimates of selection (Fig. 9; Fig. S5). The choice of stratified versus random available point positioning *β*: −0.001, 95% CrI −0.004 – 0.002 and the available points multiplier *β*: −0.002, 95% CrI −0.004 – −0.001 have less of an impact, mirroring the smaller effects seen in the absolute deviation model. All area related choices do not move relative to each other, and reveal that MCPs as an area method lead to lower estimates of selection in this scenario.

Every decision examined by the RSF absolute effect model appears to be make a difference when estimating selection (Fig. 9; Fig. S2). Like the Wides analysis, data quantity is most important. The two largest effects are from tracking duration (*β*: −0.447, 95% CrI −0.47 – −0.425) and frequency (*β*: −0.609, 95% CrI −0.631 – −0.588), showing that more data leads to more stable estimates of habitat selection (i.e., more likely to get an answer closer to the overall median estimate).

However, unlike with Wides, the other analysis decisions also matter, although to varying extents. It appears that larger contours (*β*: 0.018, 95% CrI 0.007 – 0.028) and a greater multiplier of available points (*β*: −0.071, 95% CrI −0.08 – −0.06) decreases the deviation from the median estimate. To a small extent, using a stratified available points pattern led to lower deviation from the median than a purely random available points pattern (*β*: 0.001, 95% CrI −0.019 – 0.025). Choice of area method can alter results in differing ways. Compared to using MCPs, both AKDE (*β*: 0.057, 95% CrI 0.025 – 0.086) and KDEhref (*β*: 0.041, 95% CrI 0.012 – 0.075) produce estimates more likely to be closer to the median estimate, and to similar degrees. By contrast, dynamic Brownian Bridge Movement Models are more likely to lead to greater variation in estimates than those analyses that use MCPs (*β*: 0.124, 95% CrI 0.096 – 0.154).

Examining the interactions between tracking duration and frequency with the area methods further reveals the connection between dBBMMs and more variable estimates. While the other area methods interact with tracking duration in a similar fashion, dBBMMs are more sensitive, meaning that at shorter tracking durations the variation created by choosing dBBMMs over another method is greater (Fig. 10). When interacting with tracking frequency, the sensitivity is less pronounced; instead we find MCPs being the most sensitive to a lowering of tracking frequency (Fig. 10).

**Figure 10.**
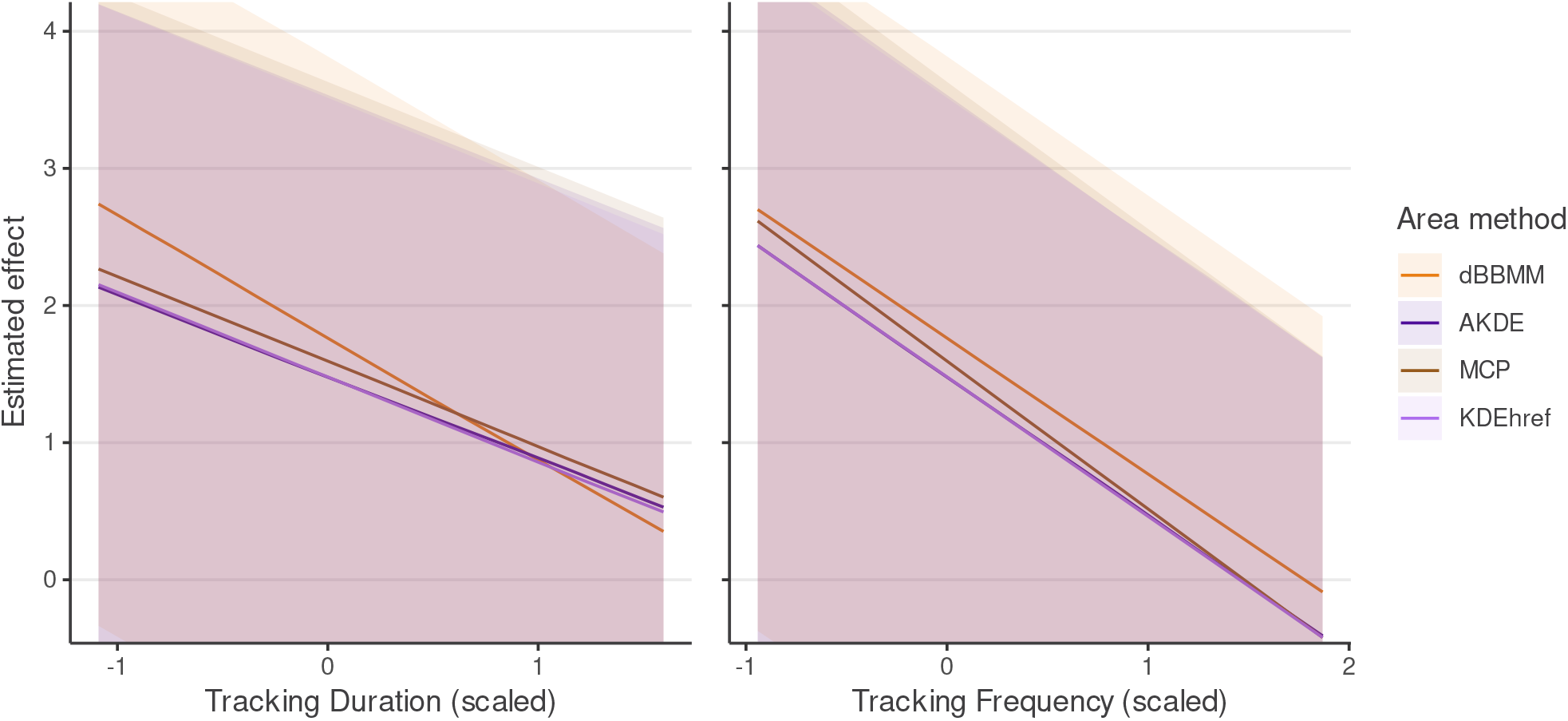
The resulting interactions from the Bayesian regression model looking to explain absolute deviation from the median in the RSF analysis pathway. Left-hand plot shows how scaled tracking duration interacted with area method chosen, while the right-hand shows the interaction between scaled tracking frequency and area method chosen.

The RSF model examining raw deviation from the median estimate reveals a general tendency for higher data quantity to lead to more conservative estimates of selection (tracking duration *β*: −0.231, 95% CrI −0.256 – −0.207 and tracking frequency *β*: −0.265, 95% CrI −0.291 – −0.243; Fig. 9; Fig. S6). While the way the movement data is simulated makes it impossible to definitively determine whether this conservatism is correct, the direct estimates from the simulations using an RSF approach suggest the low positive estimates (also near the median; see associated data for directly estimated selection values, Table. S2) are more likely to be the correct estimation of simulated selection via RSF. This matches the absolute deviation model where more data leads to less deviant estimations.

Stratified available point sampling was associated with more positive estimates of selection (*β*: 0.016, 95% CrI −0.007 – 0.038), but also those closer to the median (as indicated by the previous absolute deviation model). This suggests that random point sampling, by contrast, leads to more variable estimates that tend to be lower. The same pattern is visible in available points multiplier, where more points lead to a more consistent estimate but with a tendency for that estimate to be higher (*β*: 0.312, 95% CrI 0.301 – 0.324).

The area method choices reveal a similar pattern. Dynamic Brownian Bridge Movement models were connected to the more variable estimates, and that variability tended towards a lower estimates (*β*: 0.252, 95% CrI 0.221 – 0.286). The other methods are connected to lower deviation from the median estimate, but higher raw estimates. The area contour selected also reflects this (*β*: 0.325, 95% CrI 0.312 – 0.336). Taken together, it appears that methods capturing a spatially broader picture of availability have a tendency to produce more stable (in relation to other analysis decisions) but higher estimates of selection. The dBBMMs, by contrast, are more vulnerable to variation in other decisions, a trait reflected in the interaction with data quantity measures.

Overall there does not appear to be a decision that leads to great deviation from the median estimate in one direction.

The SSF absolute deviation model very closely matches the previous two models (Fig. 9; Fig. S3), where tracking quantity both in terms of duration (*β*: −0.742, 95% CrI −0.791 – −0.688) and frequency (*β*: −1.13, 95% CrI −1.176 – −1.079) dominate in regards to explaining why estimates tend to be closer to the median estimate. Unlike the other analysis, all other decisions made a negligible impact on an estimate’s absolute proximity to the median estimate.

Due to the one sided variability of the SSF estimates the raw deviation model mostly mirrors the absolute deviations (Fig. 9; Fig. S7). The only noticeable difference is in the model formula decision that changes the direction of the beta point estimate from negative to positive (*β*: 0.017, 95% CrI −0.076 – 0.116). However, the large overlap with zero means that in both cases the step distribution decision is not making a considerable impact to the estimates relative to the median estimate.

The results from the wRSF models are largely similar to the SSF models, although with fewer decisions (Fig. 9; Fig. S4). Both tracking duration and tracking frequency lead to lower deviation from the median estimate (tracking duration *β*: −0.308, 95% CrI −0.459 – −0.156 and tracking frequency *β*: −0.193, 95% CrI −0.352 – −0.044), and the near-exclusively positively skewed outliers mean that lower raw deviation is associated with lower estimates (tracking duration *β*: −0.3, 95% CrI −0.47 – −0.129 and tracking frequency *β*: −0.207, 95% CrI −0.383 – −0.039; Fig. 9; Fig. S4). The only difference (although both effects overlap considerably) is that increased tracking duration tended to be linked more strongly with lower selection estimates.

## 4 Discussion

Our multiverse exploration of individual habitat selection analysis methods reveals the importance of data quantity in generating consistent results. Regardless of the approach used, data quantity increases resulted in estimates of habitat selection closer to the median estimate and more consistently in line with the underlying simulated habitat selection. There appeared to be a reduction in deviation from the median estimate when using newer methods (e.g., SSF and wRSF). This suggests that advancements in habitat selection analysis methods are not only worthwhile, but actively counter reproducibility issues that may stem from researcher choice.

The overall pattern in estimates mirrors those seen in other fields, where the bulk of answers converge but outliers remain (Silberzahn et al., 2018; Simonsohn, Simmons & Nelson, 2020; Gould et al., 2023). While the incentives for findings positive results remain in the publishing system (Fanelli, 2010b, 2012), in the case presented here, using analysis choices to game or mine for a given exciting result appears time consuming because the proportion of extreme estimates is relatively low compared to the majority of estimates that agree near the median. For example, the 10 and 90% percentiles easily exclude outlying estimates, particularly those lower estimates, and remain reasonably close to the median estimates Wides selection: 10% 0.913, 90% 1.68; Wides scrambled: 10% 0.867, 90% 1.23; RSF selection: 10% −0.52, 90% 11.3; RSF scrambled: 10% −1.12, 90% 0.881; SSF selection: 10% −0.0184, 90% 15.6; SSF scrambled: 10% −0.811, 90% 0.763; wRSF selection: 10% −0.474, 90% 2.97; wRSF scrambled: 10% −2.21, 90% 1.75. If the past habitat selection studies report extreme findings, there is a general tendency for these to be paired with very wide confidence intervals; if the studies have reported the confidence intervals, this provides an additional safeguard against over interpretation of potentially incorrect extreme results.

While the goal of this study was not to review and provide direct guidance for the most effective ways of undertaking habitat selection, our findings show several key patterns that reflect previous guidance in the literature. In particular the benefits of larger datasets and the importance of defining availability (Northrup et al., 2013; Street et al., 2021); the latter is particularly key because of the potential for subjective definitions (Beyer et al., 2010). One specific example would be the tendency for larger area methods and wider contours leading to more stable (nearer the median) estimates; this mirrors previously revealed patterns connecting broader spatial scales and transferability (Paton & Matthiopoulos, 2015).

The overall pattern shows data quality supremacy, but this cannot be used as an excuse to ignore the decisions during analysis. Two aspects of the results highlight that complacency is unwise. First RSF, SSF, and wRSF approaches appear to be more consistent than the more basic and older Wides ratio based approach. Both RSF, SSF, and wRSF have improved capacity to account for the peculiarities of movement data structure (e.g., via random effects) than the Wides approach. Second is the evidence for conceptual mistakes to have impacts on final results, implying that in some cases there are *more correct* answers to certain analysis decisions. The use of dBBMMs to define availability is not strictly correct as the (confidence) area generated by dBBMMs reflects uncertainty surrounding use. This renders the down stream analysis of habitat selection a comparison of use (measured by points) to use (measured by area of uncertainty surrounding movement trajectories); as opposed to the desired use versus availability. The dBBMM results exemplify the need to maintain a connection between abstractions (e.g., an area) of the data and the intended use. A similar argument can be made for data quantity, where mismatches between frequency and desired behaviours or an inadequate tracking duration will obscure answers regardless of the analysis applied. Our simulations are diurnal and the tracking spacing was skewed towards those diurnal hours for lower tracking frequency regimes. Failure to do this, instead allowing locations to be recorded equally at night and day, would have skewed estimates towards the habitat used for shelter rather than foraging. Matching question to analysis approach remains important, but the cost of mistakes may be less dramatic than feared for general assessments of habitat selection.

Further good news is that some decisions connected to more stable estimates have very little reason not to be maximised. For example, the points multiplier or points per step decision can be maximised to whatever extent allowed by the computational resources available to the researcher. Increases in the number of available points will improve accuracy when defining available habitats; Street et al. (2021) indicate that the systematic sampling with high numbers of points is preferable to capture a complete unbiased availability. Fieberg et al. (2021) states that available points should be high enough to ensure stable coefficient estimates, that is also aided by increased weighting of available points in the logistic regression. While we did not see the improvements in points per step in the SSF analysis, the benefits of high available steps is more pronounced when working with rare habitat types (Thurfjell, Ciuti & Boyce, 2014). More available points reduces the chances of missing small areas of rare habitat; our analysis would not have revealed this because of the abundance of the preferred habitat type.

We did not see large improvements using different step or turn angle distributions; this has been shown to play an increasingly important role in mitigating biased estimates in scenarios with stronger selection strengths (Forester, Im & Rathouz, 2009). The limited cost of integrating step and turn into SSF formulas would suggest iSSF formulation is likely always preferable for estimation of individual habitat selection.

### 4.1 Limitations of the multiverse approach and future directions

While this study helps demonstrate the impact of researcher choice in habitat selection studies, the true number of choices is impractical to explore and ever growing. Time and computational limits compound with the infinite granularity of decisions connected to continuous variables (e.g., tracking frequency) to ensure that all choices cannot be completely explored.

The relationship between tracking regime, data quantity produced, and the resulting habitat selection results is a complex area to assess. We focused on systematic controlled changes to tracking frequency, but in field scenarios data quantity can be impacted by practical and logistical constraints that result in stochastic data loss. Newer methods help alleviate the problems with inconsistent tracking (Fleming et al., 2018; Silva et al., 2022; Alston et al., 2023), providing an additional reason why analysis decisions should not be ignored.

Ultimately the multiverse approach cannot reveal the true answer; the multiverse’s mean result describes the mean of the analysis pathways and does not necessarily converge towards a true effect. In much the same way as replication does not provide a direct proxy for truth, the chosen analysis pathways, and their relation to an underlying truth, is not necessarily improved by conducting more. Improvement will only come with the additions being fundamentally more accurate, and better reflective of the study system (Devezer et al., 2021).

This unclear relationship between consistency of results and the study system is why we tried to capture a range of variability in the simulations. We attempted to make our findings more generalisable by simulating three distinct animal species that move in different fashions and that face differing levels of habitat connectivity. We cannot discount scenarios with different movement characteristics that interact with particular analysis choices to bias habitat selection estimates. Scale remains a key consideration for any study (Thurfjell, Ciuti & Boyce, 2014; Northrup et al., 2022), and context-specific selection remains difficult to generalise (Avgar, Betini & Fryxell, 2020). The difficulty in generalising a “one-size fits all” approach highlights the benefits of methods that can be parametrised by the underlying characteristics of the data (e.g., wRSF Alston et al., 2023), and where the uncertainty stemming from sampling can be carried forward to the final estimates.

Despite the inherit limitations of multiverses, the exploration of analysis choice presents an important consideration for any researcher looking to know the reliability of their findings (Hoffmann et al., 2021). More formal and mathematically based estimates of reliability are available for specific approaches, for example RSFs (Street et al., 2021). These present an important component when reporting RSFs. However, they still require some level of *a priori* decision making (e.g., landscape and availability definition) (Northrup et al., 2022); therefore, researcher choice remains.

Beyond the aforementioned limitations to the multiverse approach, there is scope to expand the exploration. Our next step is to reapply the same meta-programming pipeline approach to population targeting habitat analyses. A focus on these approaches will allows us to explore the third key component of data quantity –sample size– as well as how well analyses tackle individual variability either in the analysis itself or during summarising of individual habitat selection estimates. There are already methods for assessing required sample size to detected a given selection for RSFs (Street et al., 2021).

### 4.2 Conclusions

We show, using a multiverse of analysis pathways, that data quantity is an important safeguard against extreme outlying results in individual habitat selection analyses. The indications that analysis choices are less important than data quantity improves our confidence in previously published habitat selection studies as there is no more scope to mine a desired result than in other disciplines. Newer methods of habitat selection analysis appear to reduce variation from researcher choice further, indicating that future studies may be more reproducible than those before.

## 5 Acknowledgements

This work was supported by the Natural Environment Research Council (NERC) via the IAPETUS2 Doctoral Training Partnership held by Benjamin Michael Marshall (grant reference NE/S007431/1). We would also like to thank Aubrey Alamshah for their support and manuscript review.

## 6 Software availablity

In addition to packages already mentioned in the methods we also used the following.

We used *R* v.4.2.2 (R Core Team, 2023) via *RStudio* v.2023.12.0.369 (RStudio Team, 2022). We used *here* v.1.0.1 (Müller, 2020), *readr* v.2.1.5 (Wickham, Hester & Bryan, 2023), and *qs* v.0.26.3 (Ching, 2023) to manage directory addresses and saved objects.

We used *raster* v.3.6.26 (Hijmans, 2023) and *RandomFields* v.3.3.14 (Schlather et al., 2015) to aid landscape raster creation alongside *NLMR* v.1.1.1 (Sciaini et al., 2018).

We used *ggplot2* v.3.5.1 for creating figures (Wickham, 2016), with the expansions: *patchwork* v.1.2.0 (Pedersen, 2022), *ggridges* v.0.5.6 (Wilke, 2022), *ggtext* v.0.1.2 (Wilke & Wiernik, 2022), and *ggdist* v.3.3.2 (Kay, 2023a).

We used *brms* v.2.21.0 (Bürkner, 2021) to run Bayesian models, with dianogistics generated used *bayesplot* v.1.11.1 (Gabry et al., 2019), *tidybayes* v.3.0.6 (Kay, 2023b), and *performance* v.0.11.0 (Lüdecke et al., 2021).

We used the *dplyr* v.1.1.4 (Wickham et al., 2023), *tibble* v.3.2.1 (Müller & Wickham, 2023), *reshape2* v.1.4.4 (Wickham, 2007), and *stringr* v.1.5.1 (Wickham, 2022) packages for data manipulation.

We used *sp* v.2.1.4 (Bivand, Pebesma & Gomez-Rubio, 2013), *adehabitatHR* v.0.4.21 (Calenge & Scott Fortmann-Roe, 2023), *move* v.4.2.4 (Kranstauber, Smolla & Scharf, 2023) for manipulation of spatial data and estimation of space use not otherwise mentioned in the methods.

We used *rmarkdown* v.2.27 (Xie, Allaire & Grolemund, 2018; Xie, Dervieux & Riederer, 2020; Allaire et al., 2023), *bookdown* v.0.39 (Xie, 2016, 2022), *tinytex* v.0.51 (Xie, 2019, 2023a), *kableExtra* v.1.4.0 (Zhu, 2021), and *knitr* v.1.47 (Xie, 2014, 2015, 2023b) packages to generate type-set outputs.

We generated R package citations with the aid of *grateful* v.0.2.4 (Francisco Rodríguez-Sánchez, Connor P. Jackson & Shaurita D. Hutchins, 2023).

## 7 Data availabilty

The code used to simulate the movement data, perform the multiverse, and analysis the results is available at: https://github.com/BenMMarshall/multiverseHabitat, and archived at https://doi.org/10.5281/zenodo. 12169335. These locations also include model outputs and results not otherwise displayed here.

## 8 Supplementary Material

**Table S1.** All *R*^2^ values for Bayesian regression models.

| $R^2$ | Standard error | Confidence interval | CI Lower | CI Upper | Component | Response | Model Method |
| --- | --- | --- | --- | --- | --- | --- | --- |
| 0.097 | 0.001 | 0.95 | 0.095 | 0.099 | Conditional | Absolute Deviation | rsf |
| 0.034 | 0.000 | 0.95 | 0.033 | 0.035 | Marginal | Absolute Deviation | rsf |
| 0.078 | 0.001 | 0.95 | 0.077 | 0.080 | Conditional | Raw Deviation | rsf |
| 0.029 | 0.000 | 0.95 | 0.028 | 0.030 | Marginal | Raw Deviation | rsf |
| 0.279 | 0.002 | 0.95 | 0.274 | 0.284 | Conditional | Absolute Deviation | wides |
| 0.023 | 0.001 | 0.95 | 0.021 | 0.025 | Marginal | Absolute Deviation | wides |
| 0.302 | 0.002 | 0.95 | 0.297 | 0.306 | Conditional | Raw Deviation | wides |
| 0.030 | 0.001 | 0.95 | 0.029 | 0.032 | Marginal | Raw Deviation | wides |
| 0.225 | 0.004 | 0.95 | 0.218 | 0.232 | Conditional | Absolute Deviation | ssf |
| 0.064 | 0.002 | 0.95 | 0.059 | 0.068 | Marginal | Absolute Deviation | ssf |
| 0.226 | 0.004 | 0.95 | 0.219 | 0.233 | Conditional | Raw Deviation | ssf |
| 0.062 | 0.002 | 0.95 | 0.057 | 0.066 | Marginal | Raw Deviation | ssf |
| 0.088 | 0.019 | 0.95 | 0.051 | 0.125 | Conditional | Absolute Deviation | wrsf |
| 0.021 | 0.010 | 0.95 | 0.005 | 0.043 | Marginal | Absolute Deviation | wrsf |
| 0.168 | 0.022 | 0.95 | 0.125 | 0.212 | Conditional | Raw Deviation | wrsf |
| 0.016 | 0.008 | 0.95 | 0.003 | 0.032 | Marginal | Raw Deviation | wrsf |

**Figure S1.**
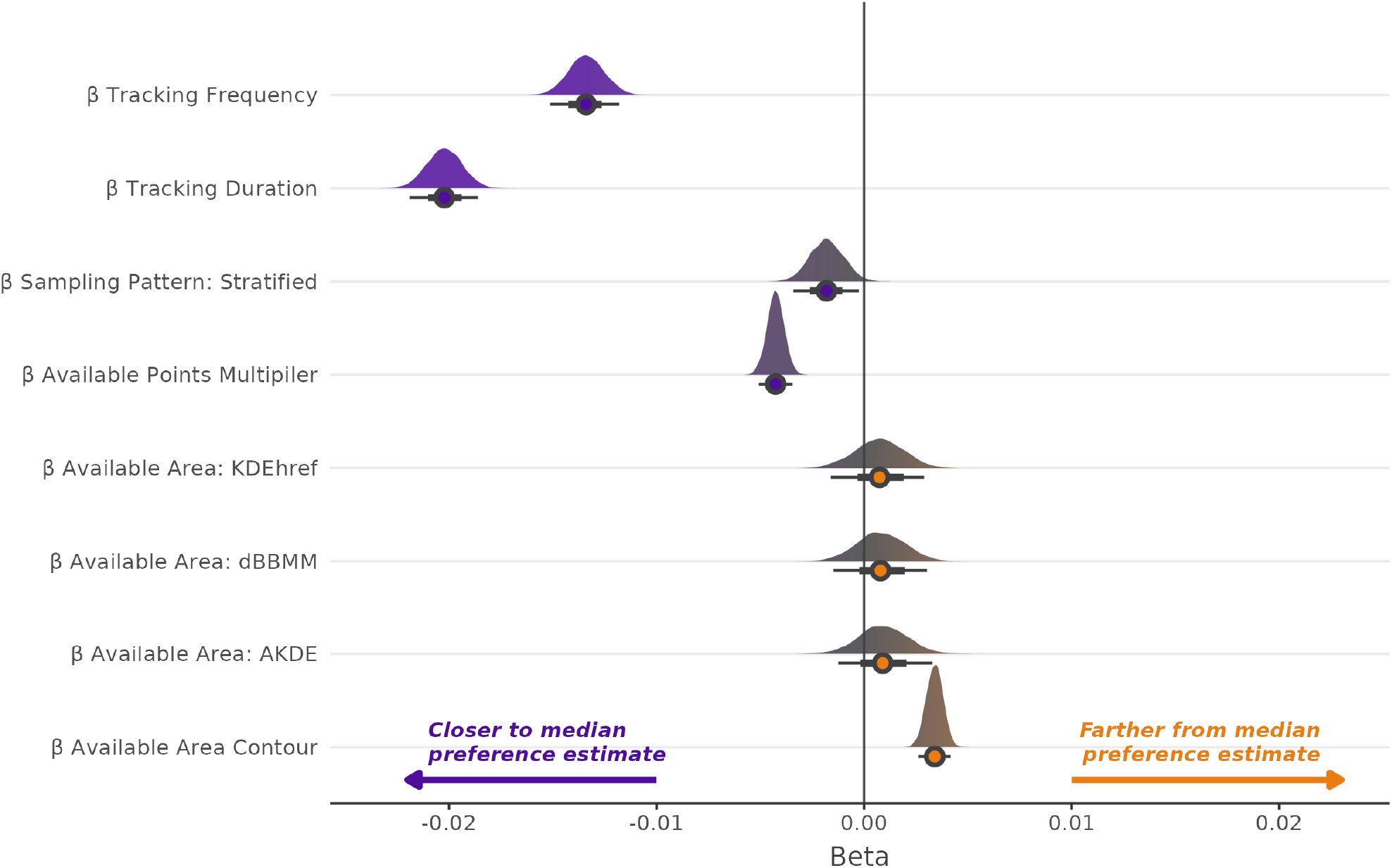
The posterior distributions of the betas from the Bayesian regression model looking to explain absolute deviation from the median in the Wides analysis pathway.

**Figure S2.**
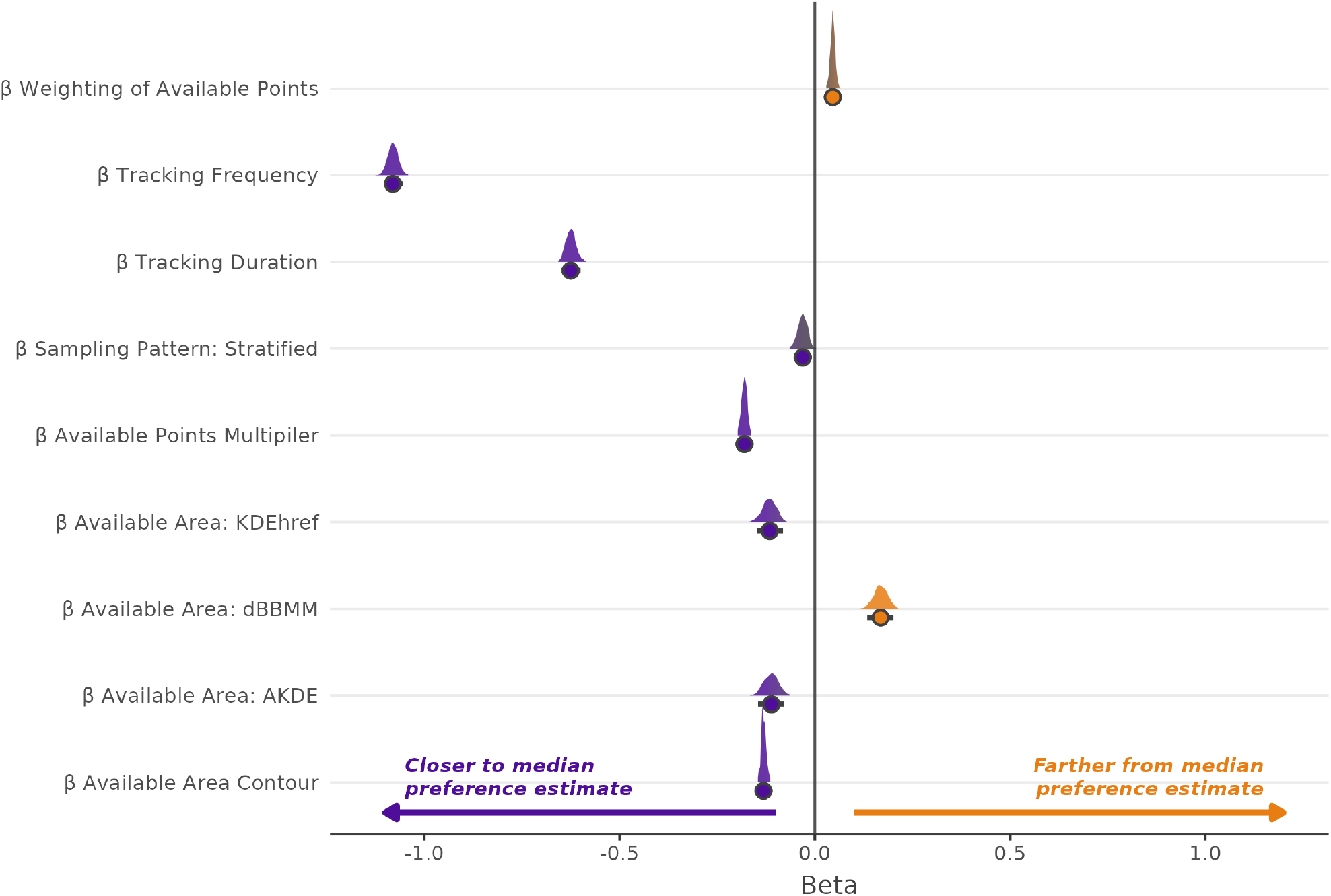
The posterior distributions of the betas from the Bayesian regression model looking to explain absolute deviation from the median in the RSF analysis pathway.

**Figure S3.**
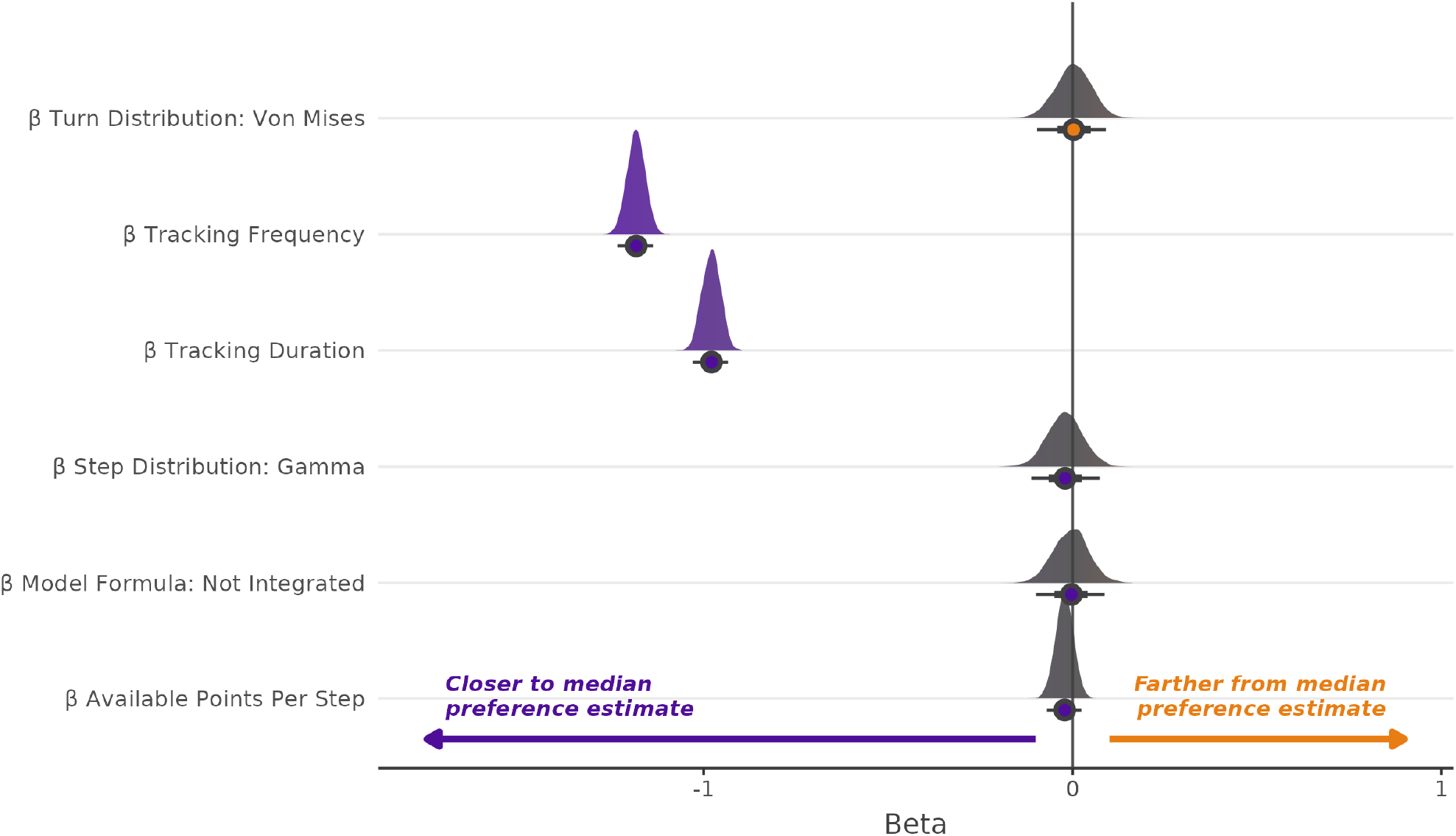
The posterior distributions of the betas from the Bayesian regression model looking to explain absolute deviation from the median in the SSF analysis pathway.

**Figure S4.**
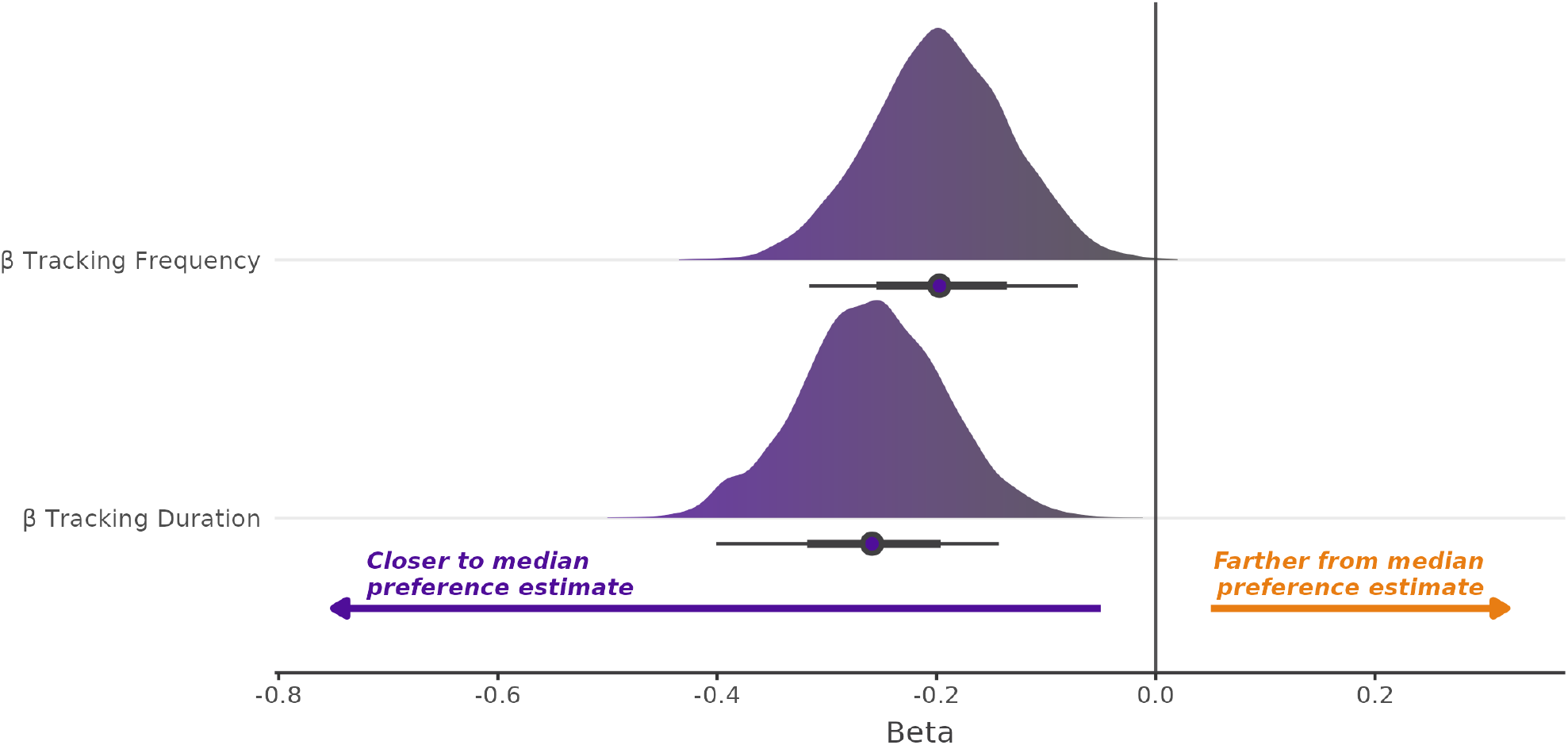
The posterior distributions of the betas from the Bayesian regression model looking to explain absolute deviation from the median in the wRSF analysis pathway.

**Figure S5.**
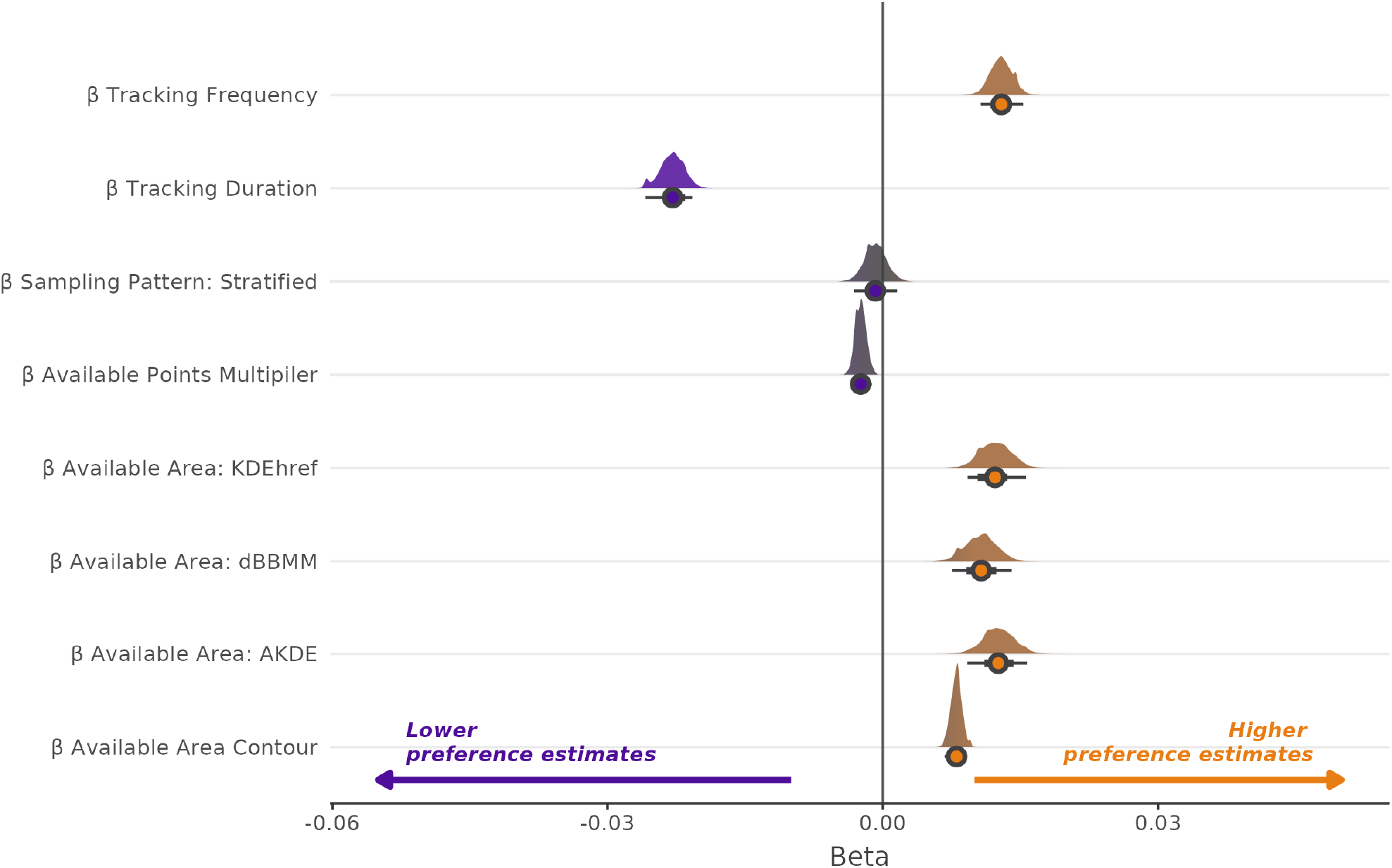
The posterior distributions of the betas from the Bayesian regression model looking to explain raw deviation from the median in the Wides analysis pathway.

**Figure S6.**
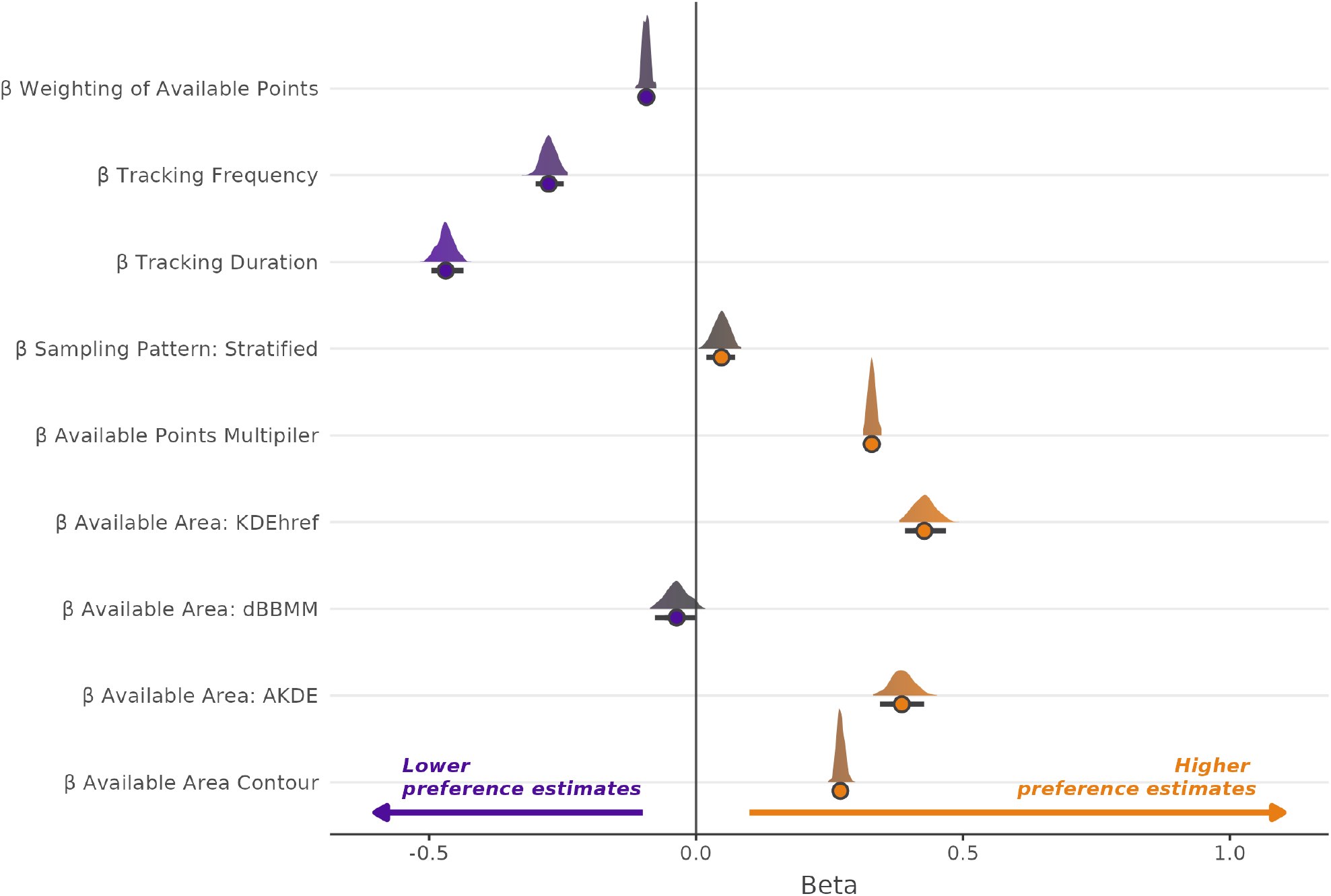
The posterior distributions of the betas from the Bayesian regression model looking to explain raw deviation from the median in the RSF analysis pathway.

**Figure S7.**
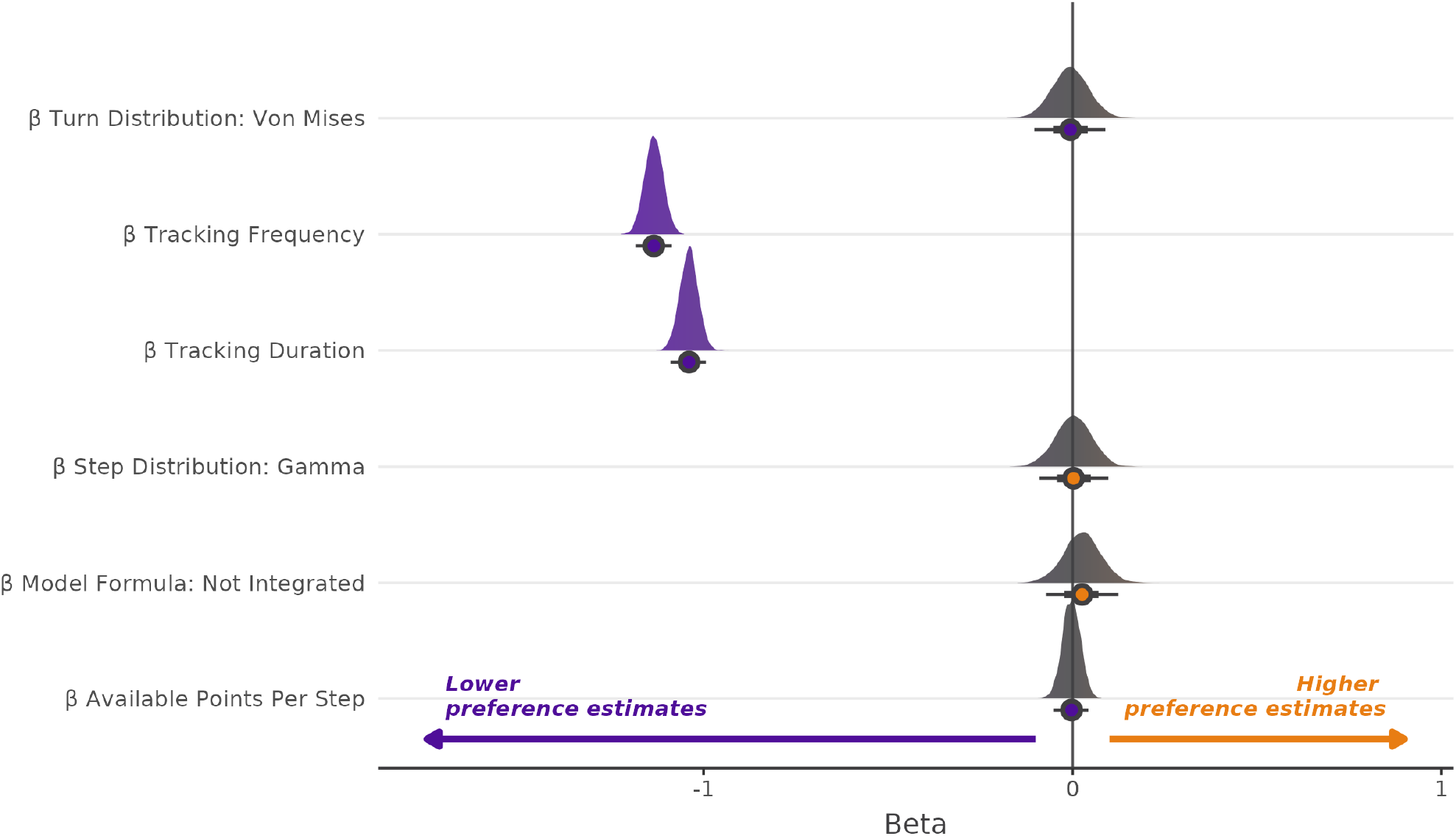
The posterior distributions of the betas from the Bayesian regression model looking to explain raw deviation from the median in the SSF analysis pathway.

**Table S2.** Analysis results when applied directly to un-resampled movement data at both the decision timescale and movement timescales.

| Estimate | Standard error | Analysis method | Simulation scale | Individual | Species |
| --- | --- | --- | --- | --- | --- |
| 0.019 | 0.059 | rsf | Movement | 1 | BADGER |
| 1.000 | NA | wides | Movement | 1 | BADGER |
| 0.078 | 0.076 | ssf | Movement | 1 | BADGER |
| 0.281 | 0.223 | rsf | Destination | 1 | BADGER |
| 1.087 | NA | wides | Destination | 1 | BADGER |
| -0.002 | 0.223 | ssf | Destination | 1 | BADGER |
| -0.002 | 0.011 | rsf | Movement | 1 | VULTURE |
| 1.002 | NA | wides | Movement | 1 | VULTURE |
| 0.005 | 0.016 | ssf | Movement | 1 | VULTURE |
| 0.344 | 0.065 | rsf | Destination | 1 | VULTURE |
| 1.202 | NA | wides | Destination | 1 | VULTURE |
| -0.114 | 0.066 | ssf | Destination | 1 | VULTURE |
| -0.009 | 0.010 | rsf | Movement | 1 | KINGCOBRA |
| 1.001 | NA | wides | Movement | 1 | KINGCOBRA |
| -0.022 | 0.017 | ssf | Movement | 1 | KINGCOBRA |
| 0.081 | 0.048 | rsf | Destination | 1 | KINGCOBRA |
| 1.082 | NA | wides | Destination | 1 | KINGCOBRA |
| -0.172 | 0.051 | ssf | Destination | 1 | KINGCOBRA |
| 0.014 | 0.008 | rsf | Movement | 2 | BADGER |
| 1.002 | NA | wides | Movement | 2 | BADGER |
| 0.046 | 0.014 | ssf | Movement | 2 | BADGER |
| 0.357 | 0.054 | rsf | Destination | 2 | BADGER |
| 1.179 | NA | wides | Destination | 2 | BADGER |
| -0.072 | 0.056 | ssf | Destination | 2 | BADGER |
| 0.004 | 0.013 | rsf | Movement | 2 | VULTURE |
| 1.006 | NA | wides | Movement | 2 | VULTURE |
| 0.027 | 0.019 | ssf | Movement | 2 | VULTURE |
| 0.293 | 0.065 | rsf | Destination | 2 | VULTURE |
| 1.158 | NA | wides | Destination | 2 | VULTURE |
| -0.183 | 0.066 | ssf | Destination | 2 | VULTURE |
| -0.001 | 0.018 | rsf | Movement | 2 | KINGCOBRA |
| 1.001 | NA | wides | Movement | 2 | KINGCOBRA |
| 0.017 | 0.028 | ssf | Movement | 2 | KINGCOBRA |
| 0.322 | 0.090 | rsf | Destination | 2 | KINGCOBRA |
| 1.057 | NA | wides | Destination | 2 | KINGCOBRA |
| 0.150 | 0.094 | ssf | Destination | 2 | KINGCOBRA |
| 0.021 | 0.017 | rsf | Movement | 3 | BADGER |
| 1.001 | NA | wides | Movement | 3 | BADGER |
| 0.027 | 0.025 | ssf | Movement | 3 | BADGER |
| 0.789 | 0.086 | rsf | Destination | 3 | BADGER |
| 1.152 | NA | wides | Destination | 3 | BADGER |
| 0.446 | 0.087 | ssf | Destination | 3 | BADGER |
| 0.000 | 0.015 | rsf | Movement | 3 | VULTURE |
| 1.002 | NA | wides | Movement | 3 | VULTURE |
| 0.001 | 0.022 | ssf | Movement | 3 | VULTURE |
| 0.306 | 0.067 | rsf | Destination | 3 | VULTURE |
| 1.173 | NA | wides | Destination | 3 | VULTURE |
| -0.181 | 0.068 | ssf | Destination | 3 | VULTURE |
| 0.011 | 0.012 | rsf | Movement | 3 | KINGCOBRA |
| 1.002 | NA | wides | Movement | 3 | KINGCOBRA |
| 0.038 | 0.019 | ssf | Movement | 3 | KINGCOBRA |
| 0.194 | 0.056 | rsf | Destination | 3 | KINGCOBRA |
| 1.094 | NA | wides | Destination | 3 | KINGCOBRA |
| -0.017 | 0.059 | ssf | Destination | 3 | KINGCOBRA |
| 0.079 | 0.061 | rsf | Movement | 4 | BADGER |
| 0.999 | NA | wides | Movement | 4 | BADGER |
| 0.124 | 0.084 | ssf | Movement | 4 | BADGER |
| -0.117 | 0.125 | rsf | Destination | 4 | BADGER |
| 1.081 | NA | wides | Destination | 4 | BADGER |
| -0.417 | 0.127 | ssf | Destination | 4 | BADGER |
| 0.007 | 0.011 | rsf | Movement | 4 | VULTURE |
| 1.007 | NA | wides | Movement | 4 | VULTURE |
| 0.032 | 0.017 | ssf | Movement | 4 | VULTURE |
| 0.348 | 0.066 | rsf | Destination | 4 | VULTURE |
| 1.186 | NA | wides | Destination | 4 | VULTURE |
| -0.139 | 0.067 | ssf | Destination | 4 | VULTURE |
| 0.004 | 0.018 | rsf | Movement | 4 | KINGCOBRA |
| 1.001 | NA | wides | Movement | 4 | KINGCOBRA |
| 0.006 | 0.028 | ssf | Movement | 4 | KINGCOBRA |
| 0.212 | 0.085 | rsf | Destination | 4 | KINGCOBRA |
| 1.069 | NA | wides | Destination | 4 | KINGCOBRA |
| 0.049 | 0.089 | ssf | Destination | 4 | KINGCOBRA |

**Figure S8.**
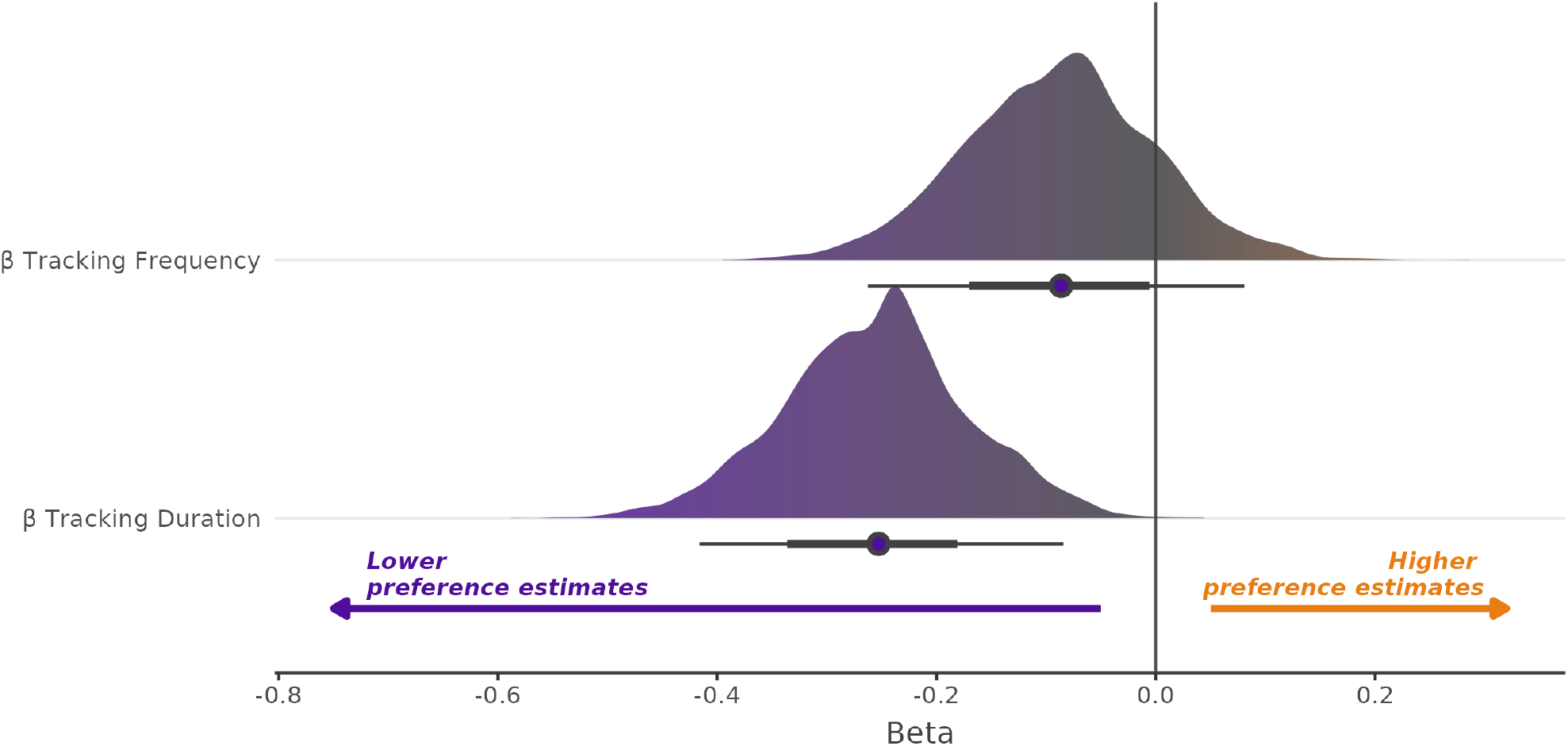
The posterior distributions of the betas from the Bayesian regression model looking to explain raw deviation from the median in the wRSF analysis pathway.

## Notes

https://zenodo.org/doi/10.5281/zenodo.12169335

### Competing Interest Statement

The authors have declared no competing interest.

